# The SMC5/6 complex prevents genotoxicity upon APOBEC3A-mediated replication stress

**DOI:** 10.1101/2023.11.28.568952

**Authors:** David R. O’Leary, Ava R. Hansen, Dylan F. Fingerman, Thi Tran, Brooke R. Harris, Katharina E. Hayer, Jiayi Fan, Emily Chen, Mithila Tennakoon, Rachel A. DeWeerd, Alice Meroni, Julia H. Szeto, Matthew D. Weitzman, Ophir Shalem, Jeffrey Bednarski, Alessandro Vindigni, Xiaolan Zhao, Abby M. Green

## Abstract

Mutational patterns caused by APOBEC3 cytidine deaminase activity are evident throughout human cancer genomes. In particular, the APOBEC3A family member is a potent genotoxin that causes substantial DNA damage in experimental systems and human tumors. However, the mechanisms that ensure genome stability in cells with active APOBEC3A are unknown. Through an unbiased genome-wide screen, we define the Structural Maintenance of Chromosomes 5/6 (SMC5/6) complex as essential for cell viability when APOBEC3A is active. We observe an absence of APOBEC3A mutagenesis in human tumors with SMC5/6 dysfunction, consistent with synthetic lethality. Cancer cells depleted of SMC5/6 incur substantial genome damage from APOBEC3A activity during DNA replication. Further, APOBEC3A activity results in replication tract lengthening which is dependent on PrimPol, consistent with re-initiation of DNA synthesis downstream of APOBEC3A-induced lesions. Loss of SMC5/6 abrogates elongated replication tracts and increases DNA breaks upon APOBEC3A activity. Our findings indicate that replication fork lengthening reflects a DNA damage response to APOBEC3A activity that promotes genome stability in an SMC5/6-dependent manner. Therefore, SMC5/6 presents a potential therapeutic vulnerability in tumors with active APOBEC3A.

## INTRODUCTION

Cytidine deamination caused by APOBEC3 enzymes is among the most prevalent sources of endogenous mutagenesis in human cancers(Nik-Zainal et al. 2012; Burns et al. 2013b; Roberts et al. 2013; Chan et al. 2015; Cortez et al. 2019; Alexandrov et al. 2020; Jalili et al. 2020). APOBEC3 enzymes catalyze the conversion of cytidine to uracil in single-stranded (ss)DNA substrates, which can result in mutations after replication or uracil excision(Chen et al. 2006; Richardson et al. 2014). The APOBEC3 enzymes function in the innate immune system to deaminate and mutate viral genomes and retroelements to restrict infection and retrotransposition(Chen et al. 2006; Richardson et al. 2014; Harris and Dudley 2015). Off-target or aberrant activity of the enzymes results in damage to the cellular genome(Landry et al. 2011; Suspene et al. 2011; Burns et al. 2013a; Green et al. 2016; Haradhvala et al. 2016; Venkatesan et al. 2021; Baker et al. 2022). Of the seven-member family (APOBEC3A-H), APOBEC3A is expressed in the nucleus and causes mutagenesis in experimental systems and human tumors, which can be genotoxic at high levels(Roberts et al. 2012; Burns et al. 2013a; Cortez et al. 2019; DeWeerd et al. 2022; Petljak et al. 2022). Mutational patterns of APOBEC3A activity are conserved across yeast and mammalian experimental models(Roberts et al. 2012; Burns et al. 2013a; Taylor et al. 2013; Chan et al. 2015; Hoopes et al. 2016; Law et al. 2020; Petljak et al. 2022).

In cancer, the genotoxic potential of APOBEC3A activity can be exploited by inhibition of the essential DNA damage responses which it activates. APOBEC3A deamination at replication forks activates replication stress responses initiated by ATR kinase signaling(Buisson et al. 2017; Green et al. 2017). Inhibition of ATR abrogates the cell cycle checkpoint, enables accumulation of mutations during DNA replication, and ultimately promotes replication catastrophe as cells move through mitosis(Buisson et al. 2017; Green et al. 2017). Cytotoxicity of APOBEC3A activity upon ATR inhibition illustrates a synthetic lethal interaction and the essential nature of DNA damage responses in tumor cells undergoing mutagenesis. We employed the synthetic lethality strategy to investigate DNA damage responses that are elicited by the activity of APOBEC3A. Using a genome-wide CRISPR-based screen, we determined that optimal function of the Structural Maintenance of Chromosomes 5/6 (SMC5/6) complex is essential in cells with active APOBEC3A.

SMC5/6 is a highly conserved eight-member complex comprised of SMC5 and SMC6 as well as six non-SMC element (NSMCE) proteins(Aragon 2018). SMC5 and SMC6 dimers form the hinge-like backbone of the complex to which other subunits attach(Alt et al. 2017; Yu et al. 2022). Similar to the related condensin and cohesin SMCs, SMC5/6 interacts with DNA to influence genome stability. Notably, SMC5/6 can bind to both ssDNA and double-stranded (ds)DNA, and can stabilize ssDNA-dsDNA junctions(Chang et al. 2022; Tanasie et al. 2022). While condensin and cohesin act in chromosome folding and segregation, the function of SMC5/6 in genome maintenance is less well-defined.

Experimental suppression of SMC5/6 as well as germline defects in SMC5/6 in human syndromes result in replication and repair defects and chromosomal aberrations(Payne et al. 2014; van der Crabben et al. 2016; Venegas et al. 2020; Grange et al. 2022; Zhu et al. 2023). Despite these data indicating an important role for the complex in genome integrity, SMC5/6 deficiency in human cancer is poorly understood.

In yeast and mammalian cells, SMC5/6 co-localizes with replication-associated proteins and nascent DNA, indicating that the complex localizes to replication structures(Ampatzidou et al. 2006; Betts Lindroos et al. 2006; Barlow et al. 2013; Alabert et al. 2014; Winczura et al. 2019). Several genome maintenance roles for SMC5/6 at replication forks have been elucidated, such as regulation of fork reversal and resolution of recombination intermediates that arise due to DNA repair at impaired replication forks(Potts and Yu 2005; Chen et al. 2009; Irmisch et al. 2009; Wu et al. 2012). Additionally, SMC5/6 localizes to natural pausing sites at centromeres, telomeres, and ribosomal DNA even in the absence of genome stress, suggesting a role for support of replication through repetitive or fragile regions(Torres-Rosell et al. 2007; Barlow et al. 2013; Menolfi et al. 2015; Peng et al. 2018; Agashe et al. 2021). While the influence of SMC5/6 on replicating DNA is established, a role for protection of forks undergoing cytidine deamination has not been defined.

In this study, we discovered a synthetic lethal interaction between APOBEC3A activity and loss of SMC5/6. By modeling SMC5/6 loss in cancer cell lines, we found that APOBEC3A activity elicited high levels of DNA breaks leading to genotoxic cell death. This synthetic lethal interaction was conserved from yeast to human tumors. In cancer cells depleted of SMC5/6, deaminase-induced DNA damage was maximal during DNA replication. Intriguingly, we found that APOBEC3A activity led to an increased length of replication forks as measured by DNA fiber imaging. The increased length was dependent on PrimPol, thus is likely due to bypass of APOBEC3A-mediated replication obstacles by repriming downstream of a DNA lesion. Interestingly, the increased fork length was also dependent on SMC5/6. We propose a model in which SMC5/6 stabilizes replication forks in cells undergoing deaminase-mediated damage. These data demonstrate a new mechanism by which genome integrity is maintained in the context of APOBEC3A activity, and reveal a synthetic lethal interaction that may provide opportunities for therapeutic targeting of SMC5/6 in cancer.

## RESULTS

### Functional genomics screen identifies SMC5/6 as essential in cells with APOBEC3A activity

To identify cellular processes that ensure genome protection from mutagenic activity of APOBEC3A, we employed a genome-wide functional screening approach. THP-1 (myeloid leukemia) cells with integrated doxycycline (dox)-inducible APOBEC3A transgene (THP1-A3A) and constitutive Cas9 transgene (**Fig S1a**) were transduced with the Brunello guide RNA (sgRNA) library which includes multiple sgRNAs for each human gene as well as non-targeting control sgRNAs(Doench et al. 2016). A 1:1 ratio of lentivirus:cell was used to allow screening for knockout of each human gene independently within a pooled population (**Fig S1b**). Following transduction, control cells (-dox) were cultured in parallel with cells induced to express APOBEC3A (+dox) for 15 days (**Fig 1a**). In both groups, cells were harvested and sgRNAs were sequenced, normalized, and analyzed by three independent pipelines to generate a gene score for each gene represented in the Brunello library (**Fig S1c**). Sequencing coverage of the entire library was similar across samples, regardless of dox treatment (**Fig S1d-e**).

**Figure 1.**
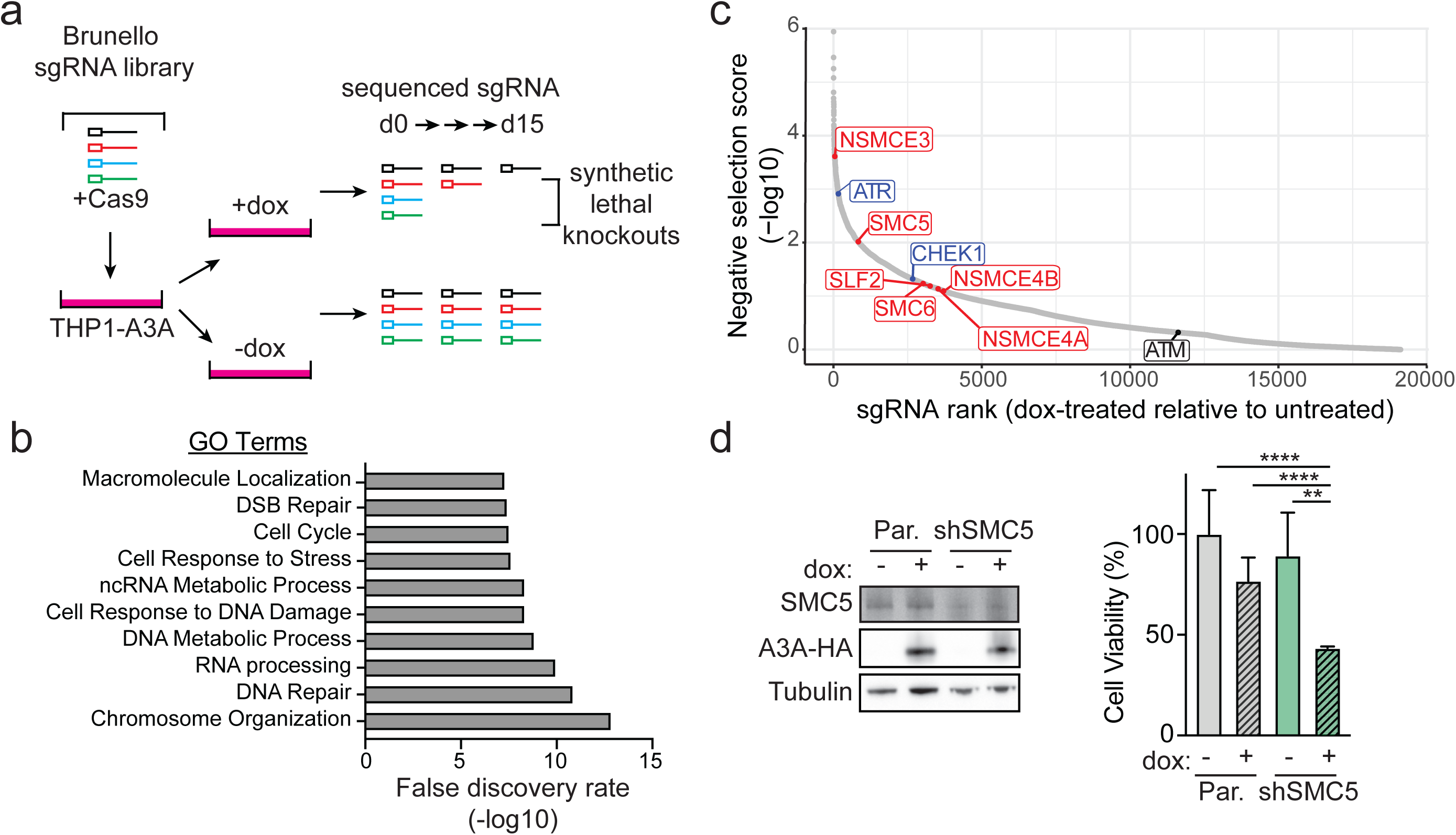
Functional genomics screen identifies synthetic lethality between loss of the SMC5/6 complex and expression of APOBEC3A. (**a**) Schematic for functional genomics screen to identify synthetic lethality with APOBEC3A. The Brunello CRISPR-Cas9 guide RNA (sgRNA) library was used in THP-1 cells expressing a doxycycline (dox)-inducible, HA-tagged APOBEC3A transgene (THP1-A3A). sgRNAs were identified and quantified by sequencing at day 0 (baseline library integration) and day 15 after dox treatment. Depletion of sgRNAs at day 15 in dox-treated cells relative to untreated controls represent potential synthetic lethal genes. (**b**) The top 250 genes identified as potentially synthetic lethal with APOBEC3A are grouped by Gene Ontology (GO) terms. (**c**) Negatively selected sgRNAs in dox-treated relative to untreated cells at day 15. SMC5/6 complex genes in red. Previously defined synthetic lethal interactions denoted in blue. (**d**) THP1-A3A cells were depleted of SMC5 by stable integration of shRNA. Viability of cells treated with dox for 72 hours or untreated was determined by FACS after staining for fluorescent-labeled calcein AM (live) and DNA (dead). Mean and SD of triplicate experiments are shown. p-values by two-tailed t-test. ****p<0.0001 **p<0.01. Cell lysates were probed with antibodies to HA and SMC5. Tubulin is loading control.

Comparison of APOBEC3A-expressing to control cells revealed under-represented or absent sgRNAs, indicating genes that were negatively selected (**Supplemental Table 1**). Negative selection is interpreted as cell death due to synthetic lethality between target gene loss and APOBEC3A expression. The top 250 negatively selected genes were analyzed for gene ontology which revealed biological processes clustered around DNA damage and repair, DNA and RNA metabolism, and chromosome organization (**Fig 1b**). Among the negatively selected genes were ATR and CHEK1 (**Fig 1c**), which have previously been demonstrated to be synthetically lethal with APOBEC3A expression(Buisson et al. 2017; Green et al. 2017). Furthermore, ATM sgRNA levels were unchanged indicating no impact on survival of APOBEC3A-expressing cells, consistent with prior findings(Buisson et al. 2017; Green et al. 2017) (**Fig 1c**).

Within chromosome organization, which was the most significant GO term, SMC5 and NSMCE3, two members of the eight-protein complex comprising SMC5/6, were significantly negatively selected (**Fig 1c**). Additional SMC5/6 genes were negatively selected, although they appeared lower on the list. To validate this potential synthetic lethal interaction, we depleted SMC5 in THP1-A3A cells and found that, upon APOBEC3A expression, cells were significantly less viable than controls (**Fig 1d**). Thus, a functional SMC5/6 complex is essential for viability of cells expressing APOBEC3A.

### SMC5/6 loss potentiates APOBEC3A-mediated genotoxicity

To explore the reproducibility of a synthetic lethal interaction between APOBEC3A and loss of SMC5/6, we depleted SMC5, SMC6, and/or NSMCE4 in cell lines from different tissues. Complex formation relies on all subunits being intact, thus depletion of one gene results in SMC5/6 dysfunction(Potts and Yu 2005; Gallego-Paez et al. 2014; Venegas et al. 2020). Prior studies demonstrated that Smc5/6 is essential in budding and fission yeast(Lehmann 2005), and complete deletion of SMC5/6 genes is embryonically lethal in mice(Ju et al. 2013), but conditional depletion of SMC5/6 is tolerated in mammalian cells(Atkins et al. 2020; Venegas et al. 2020). Thus, we used partial and inducible depletion of SMC5/6 complex genes. In K562 (myeloid) and Jurkat (T-cell) leukemia cells, shRNA was used to constitutively deplete SMC5 (**Fig 2a** and **Fig S2a**). The HCT116 colorectal carcinoma cells were engineered with auxin-inducible degron (mAID) tags on NSMCE4A and SMC6 subunits for inducible depletion of SMC5/6 upon treatment with indole-3-acetic-acid (IAA) as previously described (**Fig S2b**)(Natsume et al. 2016; Venegas et al. 2020). K562, Jurkat, and HCT116 cells were engineered to express dox-inducible APOBEC3A transgenes. The doxycycline dose used induced a level of APOBEC3A expression that resulted in minimal DNA damage. However, combined SMC5/6 depletion and APOBEC3A expression significantly impaired proliferation (**Fig 2b** and **Fig S2c-d**). Additionally, DNA damage response signaling significantly increased upon APOBEC3A expression in cells depleted of SMC5 as detected by increased phosphorylation of histone variant H2AX (gH2AX), a response to DNA breaks (**Fig 2c, Fig S2e, Fig S2g**). Consistent with these results, we found increased double-stranded DNA breaks (DSBs) by neutral comet assay (**Fig 2d** and **Fig S2f**). We hypothesized that increased DNA damage would cause cell death. Indeed, the culmination of genotoxicity was reflected by decreased viability of cells with concurrent expression of APOBEC3A and depletion of SMC5/6 (**Fig 2e** and **Fig S2h**).

**Figure 2.**
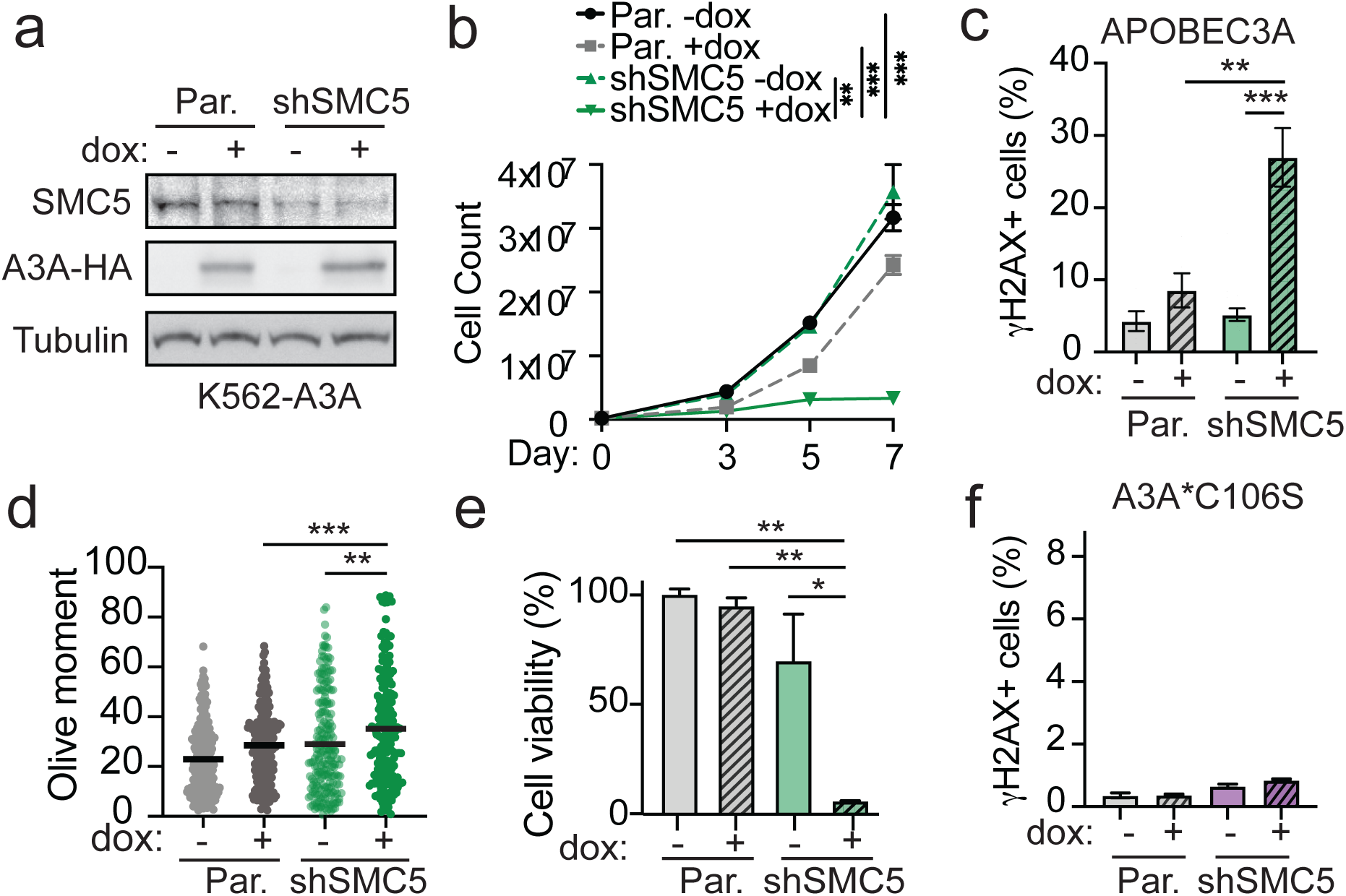
SMC5/6 loss potentiates APOBEC3A-mediated genotoxicity. K562 cells engineered to express doxycycline-inducible HA-tagged APOBEC3A (K562-A3A) were depleted of SMC5 by RNAi (shSMC5) and compared to parental K562-A3A cells. All results are representative of three independent biological replicates. (**a**) Immunoblot shows APOBEC3A expression (HA antibody) and SMC5 depletion. Tubulin was used as a loading control. (**b**) Cell proliferation was measured by counting cells over the course of 7 days. Bars are SEM, p-value by sum-of-squares F-test. (**c**) DNA damage response signaling was assessed after 72 hours of dox treatment by intracellular staining and flow cytometry analysis of the phosphorylated form of the histone variant H2AX (γH2AX). Mean and SD of triplicate experiments are shown. (**d**) Comet assay results shown as dot plot of individual values, bar is median of olive moments. (**e**) Cell viability was assessed by WST8 staining of K562 cells after 7 days of dox treatment. Mean and SD of triplicate experiments are shown. (**f**) Intracellular staining and flow cytometry analysis of γH2AX in K562 cells induced with dox for 72 hours to express the catalytically inactive A3A*C106S mutant. Mean and SD of triplicate experiments are shown. p-values by two-tailed t-test. ***p<0.001 **p<0.01 *p<0.05

### APOBEC3A catalytic activity is required for synthetic lethality with SMC5/6 loss

Next we addressed whether the synthetic lethal phenotype resulting from combined APOBEC3A expression and SMC5/6 loss was due to deaminase-induced genotoxicity. To evaluate the requirement for deamination activity, we constructed K562 cells expressing a catalytically inactive mutant of APOBEC3A containing a C106S amino acid change (**Fig S3a-b**). Cells expressing APOBEC3A-C106S with SMC5/6 depletion had no differences in proliferation, γH2AX levels, or DSB quantity (**Fig 2f** and **Fig S3c-d**). Importantly, SMC5/6 depletion did not cause increased APOBEC3A deaminase activity (**Fig S3a**). These data demonstrate that deaminase activity is required for the synthetic lethal interaction between APOBEC3A expression and SMC5/6 loss.

### SMC5/6 dsDNA binding activity protects cells from APOBEC3A toxicity in yeast

Few studies have addressed the specific functions of SMC5/6 in mammalian cells due to a lack of characterization of mutants that perturb distinct activities of the complex. Recent structural studies of the yeast Smc5/6 have enabled the generation of separation-of-function alleles of the complex that impair dsDNA binding activities(Yu et al. 2022). Given that previous studies have established yeast as a model system for studying APOBEC3A activity(Chan et al. 2015; Hoopes et al. 2016; Elango et al. 2019), we asked whether the Smc5/6 protective role against APOBEC3A toxicity is conserved in yeast, and whether its DNA binding activity is required for this protection.

A cryo-EM structure of dsDNA-bound yeast Smc5/6 complex has identified DNA binding sites on multiple subunits, with several points of contact between each subunit and DNA(Yu et al. 2022). Mutating these sites on the Smc5 and Nse4 subunits led to reduced Smc5/6 chromosome association and extreme sensitivity toward the alkylating agent MMS, suggesting that these mutations may impede DNA replication and repair needed for surviving alkylation damage(Yu et al. 2022). We thus examined whether the dsDNA binding mutant allele of Smc5 (*smc5-DNAm*, K89, K97, K98, K145, R146, R147, K192 all to A) or Nse4 (*nse4-DNAm*; R251, R256, R257, R258 all to E) were sensitive to the expression of human APOBEC3A. To do so, we transfected human APOBEC3A under a yeast promotor into mutant or wild-type cells. We confirmed comparable APOBEC3A expression in all cells (**Fig 3a-b**). Consistent with prior studies, wild-type cells were tolerant of deaminase activity as they grew similarly to those transfected with empty vector at both 30°C and 37°C (**Fig 3c-d**)(Hoopes et al. 2016). In striking contrast, yeast harboring *smc5-DNAm* or *nse4-DNAm* alleles exhibited poor viability upon APOBEC3A expression (**Fig 3c-d**). These data demonstrate that a fundamental feature of SMC5/6 in binding dsDNA that is conserved from yeast to human is responsible for supporting cell viability when APOBEC3A is active.

**Figure 3.**
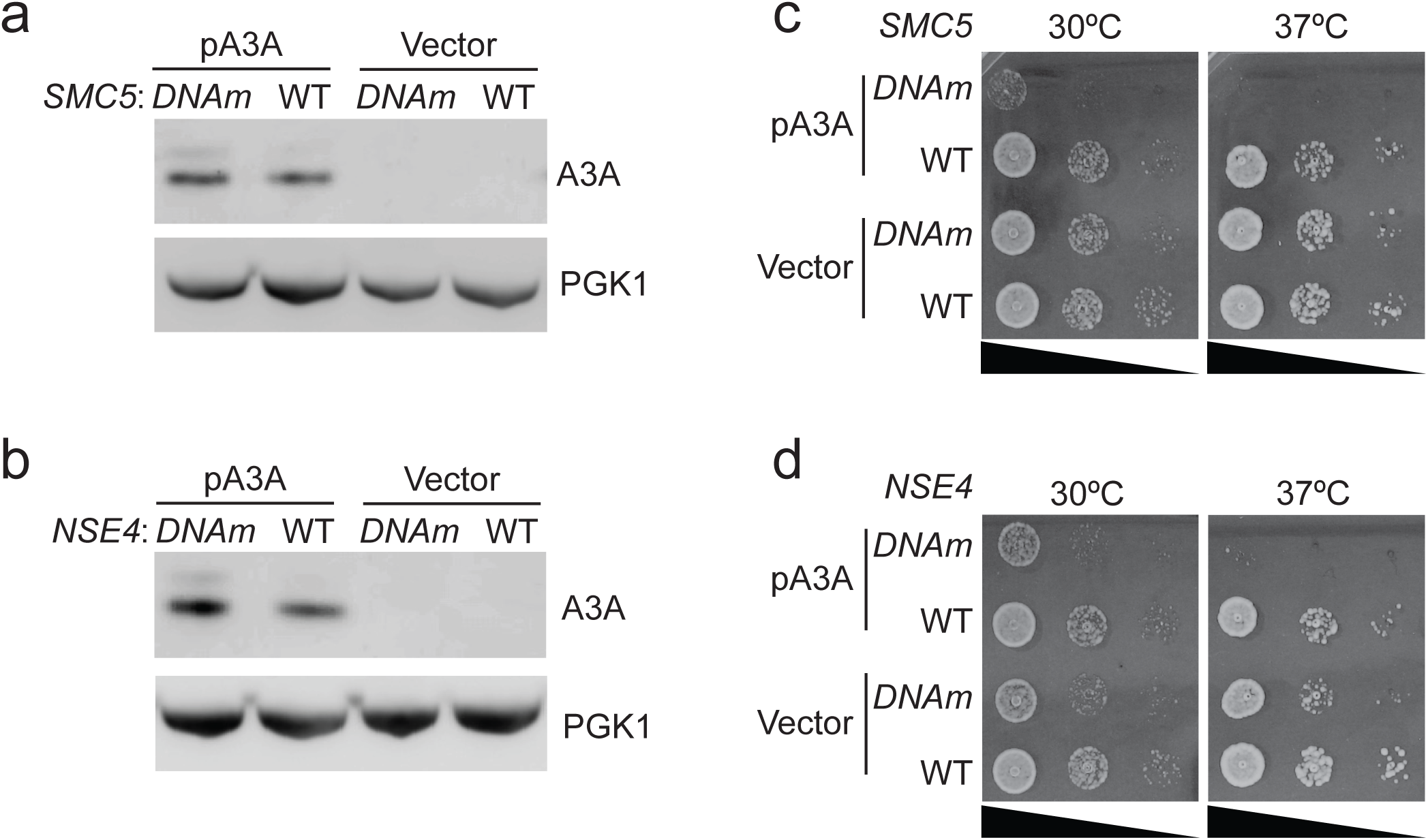
The DNA binding function of SMC5/6 is essential in yeast cells that express APOBEC3A. Wild-type (WT) and mutant yeast cells containing mutations in the DNA binding sites of Smc5 or Nse4 (*DNAm*) were examined. Cells were transformed with APOBEC3A expression plasmid (pA3A) or control vector. (**a-b**) APOBEC3A protein levels from extracts of the indicated cells were assessed by immunoblot using an antibody specific to APOBEC3A. PGK1 was used as a loading control. (**c-d**) Cells with indicated genotypes containing either vector or pA3A were analyzed for growth by spotting 10-fold serial dilutions of cells. Plates were grown for 24h at indicated temperatures.

### APOBEC3A mutagenesis is incompatible with SMC5/6 dysfunction in cancer

To evaluate the interaction of APOBEC3A activity and SMC5/6 loss in human cancers, we quantified APOBEC3A mutational signatures in tumors with deleterious mutations in SMC5/6 subunit genes. Deleterious mutations were defined as exonic missense, nonsense, or frameshift base changes(Choi et al. 2012; McLaren et al. 2016). Within TCGA, 160 tumors with deleterious mutations in SMC5/6 genes were identified (**Fig 4a**). SMC5 and SMC6 were the most frequently mutated genes of all subunits (**Fig 4b**). For comparison, we defined a control set of tumors with no mutations in SMC5/6 subunit genes (n=131) that were tissue-matched (**Fig 4a** and **Fig 4c**). Tumors with dysfunctional SMC5/6 had a higher overall mutation burden (**Fig 4d**), consistent with the role of the complex in genome integrity. To determine the source of mutagenesis in tumors with dysfunctional SMC5/6, we examined single base substitution (SBS) signatures. All SBS signatures that comprised more than 4% contribution to mutation burden within each group of tumors are shown (**Fig 4e**). The APOBEC3A signatures, SBS2 and SBS13, comprised a substantial portion of mutations in the control tumors but were notably absent in the SMC5/6-mutant tumors (**Fig 4e**). These data support our experimental findings that combined dysfunction of SMC5/6 and active APOBEC3A are incompatible in viable human tumors.

**Figure 4.**
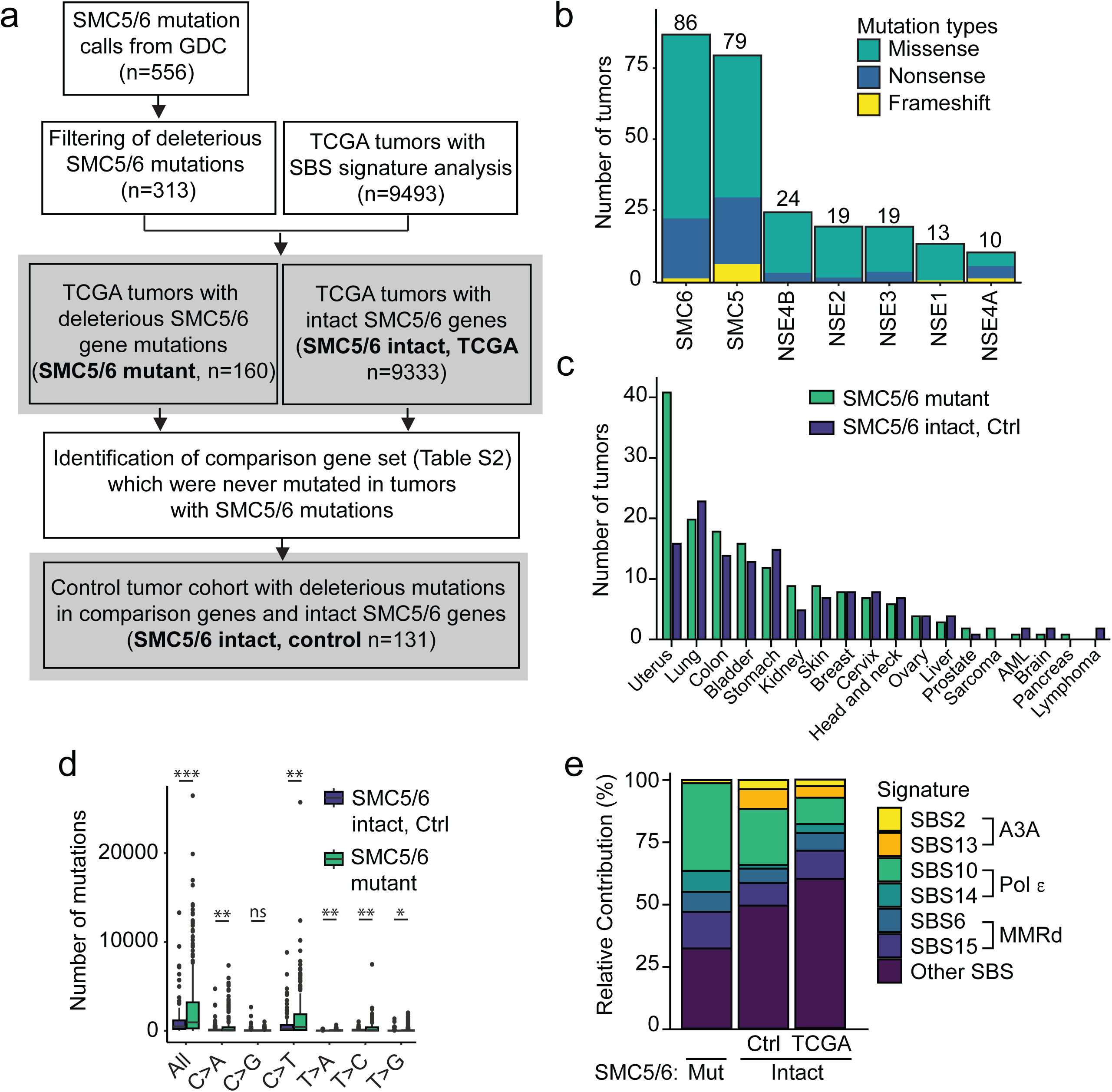
APOBEC3 mutational signatures are absent in tumors with dysfunctional SMC5/6. **(a)** Pipeline for mutational signature analysis in tumors with SMC5/6 dysfunction. Tumor genomes in the GDC data portal were evaluated for missense, nonsense, or frameshift mutations in the coding regions of SMC5/6 complex genes which would predict dysfunction. After matching with tumors in TCGA in which SBS signatures were defined, 160 tumors were identified as having mutant SMC5/6 genes. Control genes (n=40, listed in **Supplemental Table 2**) were defined by those which were never mutated in tumors with deleterious mutations in SMC5/6 genes. Tumors in which deleterious mutations in control genes were identified constituted a control (Ctrl) cohort. Bold text in gray squares indicate tumor groups included in panels **b-f**. (**b**) The number of tumors with mutations in each SMC5/6 subunit gene. (**c**) The type of tumors represented in both SMC5/6 mutant and SMC5/6 intact control groups. (**d**) Total mutation burden in each tumor genome from SMC5/6 mutant and SMC5/6 intact control cohorts. Bar is median. p-value by two-tailed t-test, *p<0.05, **p<0.01, ***p<0.001 (**e**) The relative contribution of each single base substitution (SBS) mutational signature (COSMIC v3.2) identified within the SMC5/6 mutant (Mut) and SMC5/6 intact cohorts. The latter is divided by control cohort and all other tumor genomes within TCGA. SBS signatures that comprise >4% relative contribution to mutations within each cohort are included, along with their proposed etiology. Pol ε, polymerase epsilon; MMRd, mismatch repair deficiency.

### SMC5/6 loss promotes APOBEC3A-mediated DNA damage during replication

It was previously shown that APOBEC3A activity at replication forks results in mutations on both leading and lagging strands, stalled DNA replication, and activation of DNA damage signaling(Landry et al. 2011; Green et al. 2016; Haradhvala et al. 2016; Hoopes et al. 2016; Seplyarskiy et al. 2016; DeWeerd et al. 2022). Damaged replication forks can result in replication stress and/or DNA breaks. We hypothesized that APOBEC3A activity at replication forks was a source of genotoxicity in SMC5/6-depleted cells. We used immunofluorescent staining of cyclin A to mark replicating K562-A3A cells(Sobczak-Thepot et al. 1993) and γH2AX foci to quantify DNA damage (**Fig 5a, Fig S4a**). In cells with intact SMC5/6, most APOBEC3A-induced γH2AX foci occurred in replicating cells (**Fig 5b-c, Fig S4b**). Depletion of SMC5 resulted in increased DNA damage upon APOBEC3A expression as shown by a significant increase in cells with ≥5 γH2AX foci, nearly all of which occurred in cyclin A positive cells (**Fig 5b-c, Fig S4b**). We then used a double thymidine block to synchronize cells at the G1-S junction and followed cells after release for 24 hours throughout the cell cycle (**Fig 5d**). We observed a significant accumulation of γH2AX in cells expressing APOBEC3A as they progressed through DNA replication (**Fig 5e**). Notably, APOBEC3A-expressing cells depleted of SMC5 accumulated higher levels of γH2AX throughout DNA replication relative to those with intact SMC5 (**Fig 5e**). These findings are consistent with prior reports of APOBEC3A causing genome damage during DNA replication(Green et al. 2016; Hoopes et al. 2016; Seplyarskiy et al. 2016), which we now show is exacerbated by loss of SMC5/6.

**Figure 5.**
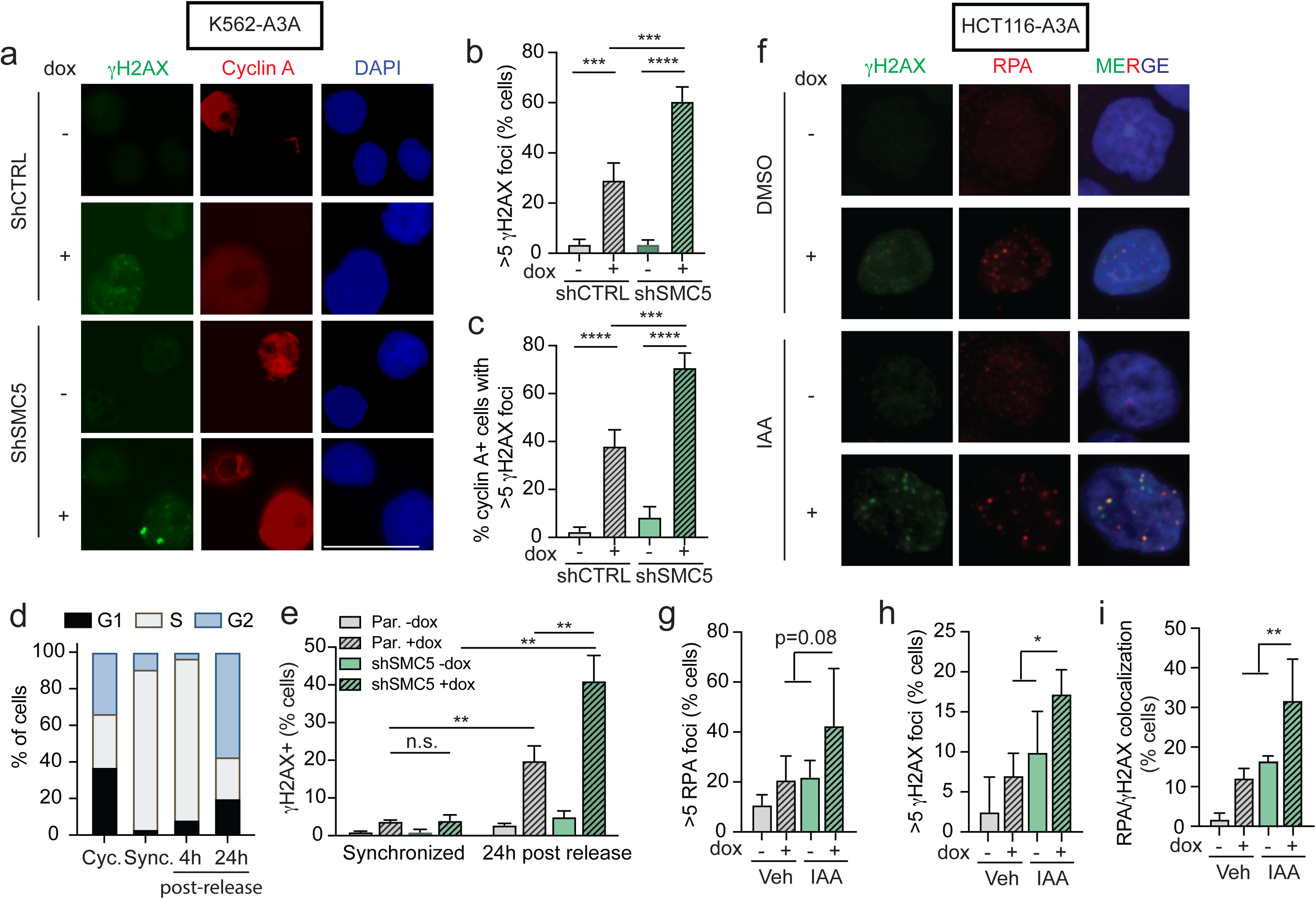
DNA damage caused by the combination of APOBEC3A activity and SMC5/6 loss occurs during DNA replication. (**a-c**) K562-A3A cells were treated with dox for 72 hours then analyzed by immunofluorescent staining of cyclin A and γH2AX. (**a**) Representative images are shown. Scale bar is 25μm. DAPI stains nuclei. (**b**) Quantification of nuclei with ≥5 γH2AX foci. (**c**) Quantification of cyclin A-positive cells that had ≥5 nuclear γH2AX foci. (**b-c**) Error bars are SEM, p-value by two-tailed t-test. (**d**) K562-A3A cells were synchronized by double thymidine block (sync.) then released. Dox was added after first thymidine block. Cells were collected 4 hours and 24 hours after release and compared to asynchronously cycling (cyc.) cells. Cell cycle was analyzed by propidium iodide (PI) staining. Bars are mean of three biological replicates. (**e**) Cells were analyzed for intracellular γH2AX staining by flow cytometry after synchronization and release. Error bars are SD, n=3, p-value by two-tailed t-test. (**f-i**) HCT116 cells with integrated E3 OsTIR1 ligase and mAID-tagged NSE4A and SMC6 subunits were treated with IAA to degrade SMC5/6 components and with dox to induce APOBEC3A expression for 72 hours. (**f**) Representative images showing RPA foci, γH2AX foci, and DAPI staining (blue). (**g-i**) Quantification of cells with >5 RPA foci (**g**), >5 γH2AX foci (**h**), and co-localized foci (**i**). Bars are mean with SD of 3 biological replicates. At least 200 nuclei were analyzed per condition. p-value by nested Anova. ****p<0.001, ***p<0.001, **p<0.01, *p<0.05.

Circumstances in which replication forks are stalled may provide more ssDNA substrate for deamination events. SMC5/6 has been implicated in perturbations and control of the G2/M cell cycle checkpoint. In plants, defective SMC5/6 promotes cell cycle progression upon DNA damage despite appropriate activation of the replication checkpoint(Wang et al. 2018). In yeast, Nse2-mediated SUMOylation of Rqh1, a RecQ helicase, is important for replication checkpoint signaling(Khan et al. 2022). Therefore, we evaluated cell cycle profiles in SMC5/6-depleted cells to determine whether replication fork stalling could explain the exacerbated genotoxicity from APOBEC3A activity. We found that cell cycle profiles were unchanged by SMC5/6 loss (**Fig S4c**). Given the exclusive activity of APOBEC3A on ssDNA substrates (as compared to dsDNA), we evaluated availability of ssDNA in cells depleted of SMC5/6. We observed that SMC5 depletion did not alter the amount of nascent ssDNA as detected by native BrdU staining (**Fig S4d**). These data show that neither cell cycle perturbations nor ssDNA substrate availability explain the excessive genotoxicity caused by APOBEC3A in the absence of SMC5/6.

We next examined localization of γH2AX foci with respect to sites of replication stress labeled by RPA foci. Following combined SMC5/6 depletion and APOBEC3A induction in HCT116-A3A cells, we observed an increase in RPA and γH2AX foci relative to controls (**Fig 5f-h**). A substantial increase in co-localization of RPA and γH2AX foci was detected in cells with both APOBEC3A expression and SMC5/6 depletion (**Fig 5i**). These data demonstrate a physical proximity of replication stress and DSB signaling, which suggests that DNA breaks may be arising from damaged replication forks.

### SMC5/6 is required for replication tract lengthening in APOBEC3A-expressing cells

To understand how replication forks were affected by deaminase activity upon SMC5/6 loss, we examined the impact of APOBEC3A activity on replication fork dynamics using single-molecule DNA fiber spreading(Quinet et al. 2017). Cells were pulsed sequentially with thymidine analogues IdU (red) and CldU (green) for a duration of 30 minutes each then analyzed for total replication tract length (IdU + CldU) (**Fig 6a**). Surprisingly, we found that APOBEC3A expression resulted in a dose-dependent increase in total tract length, indicative of replication fork elongation (**Fig 6b**). APOBEC3A-mediated fork elongation was observed in multiple cell types (**Fig 6c-f**) and was dependent on deaminase activity (**Fig 6g-h**). In all cell types, SMC5/6 depletion led to abrogation of APOBEC3A-dependent replication fork lengthening (**Fig 6c-f**). Interestingly, SMC5/6 depletion mitigated the fork elongation caused by APOBEC3A yet also resulted in DNA damage and cell death (**Fig 2**). Together, these data suggest that the activity of SMC5/6 which enables replication elongation in the context of APOBEC3A activity is protective against genotoxicity.

**Figure 6.**
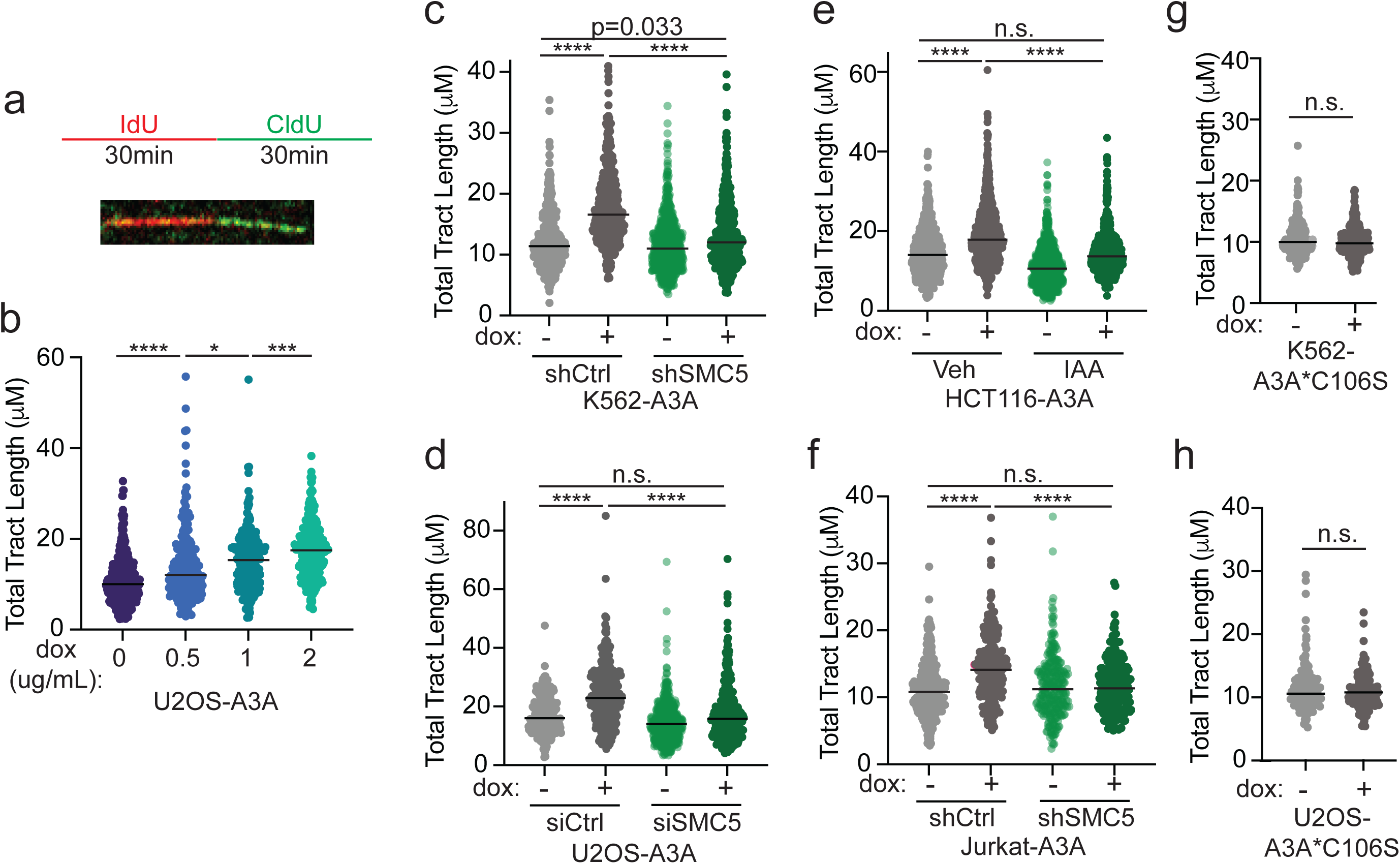
SMC5/6 is required for APOBEC3A-mediated replication fork elongation. (**a**) Schematic of DNA fiber assay and representative fiber tract. (**b**) Total tract length (μM) of DNA fibers (CldU+IdU) representing complete replication tracts from U2OS-A3A cells treated with indicated dox doses for 72 hours. (**c-f**) Total tract length (μM) of complete fibers (CldU+IdU) in K562-A3A (**c**), USOS-A3A (**d**), HCT116-A3A (**e**), and Jurkat-A3A cells (**f**) treated with dox for 72 hours. (**g-h**) Total tract length (μM) of DNA fibers from K562 (**g**) and U2OS (**h**) cells induced with dox for 72 hours to express catalytically inactive APOBEC3A (A3A*C106S). DNA fiber assays were performed in biological triplicate and analyzed by Kruskal-Wallis test. Bars are median. ****p<0.0001, ***p<0.001, *p<0.05.

### PrimPol promotes APOBEC3A-mediated elongation of replication tracts

Given the unexpected finding of increased replication tract length upon deaminase activity, we sought to determine the mechanism by which APOBEC3A leads to longer replication tracts. APOBEC3A catalyzes the conversion of cytidine to uracil which is excised by DNA glycosylases, leaving an abasic site(Chen et al. 2006; Richardson et al. 2014) which presents an obstacle for replicative polymerases. The dual primase-polymerase, PrimPol, is capable of re-initiating DNA synthesis downstream of replication obstacles(Mouron et al. 2013; Quinet et al. 2020; Quinet et al. 2021). To determine if PrimPol functions in repriming downstream of APOBEC3A-mediated DNA lesions, we expressed APOBEC3A or empty vector (EV) in PrimPol-knockout U2OS cells(Quinet et al. 2020) and measured replication tract length. Loss of PrimPol abrogated the replication tract lengthening generated by expression of APOBEC3A (**Fig 7a**).

**Figure 7.**
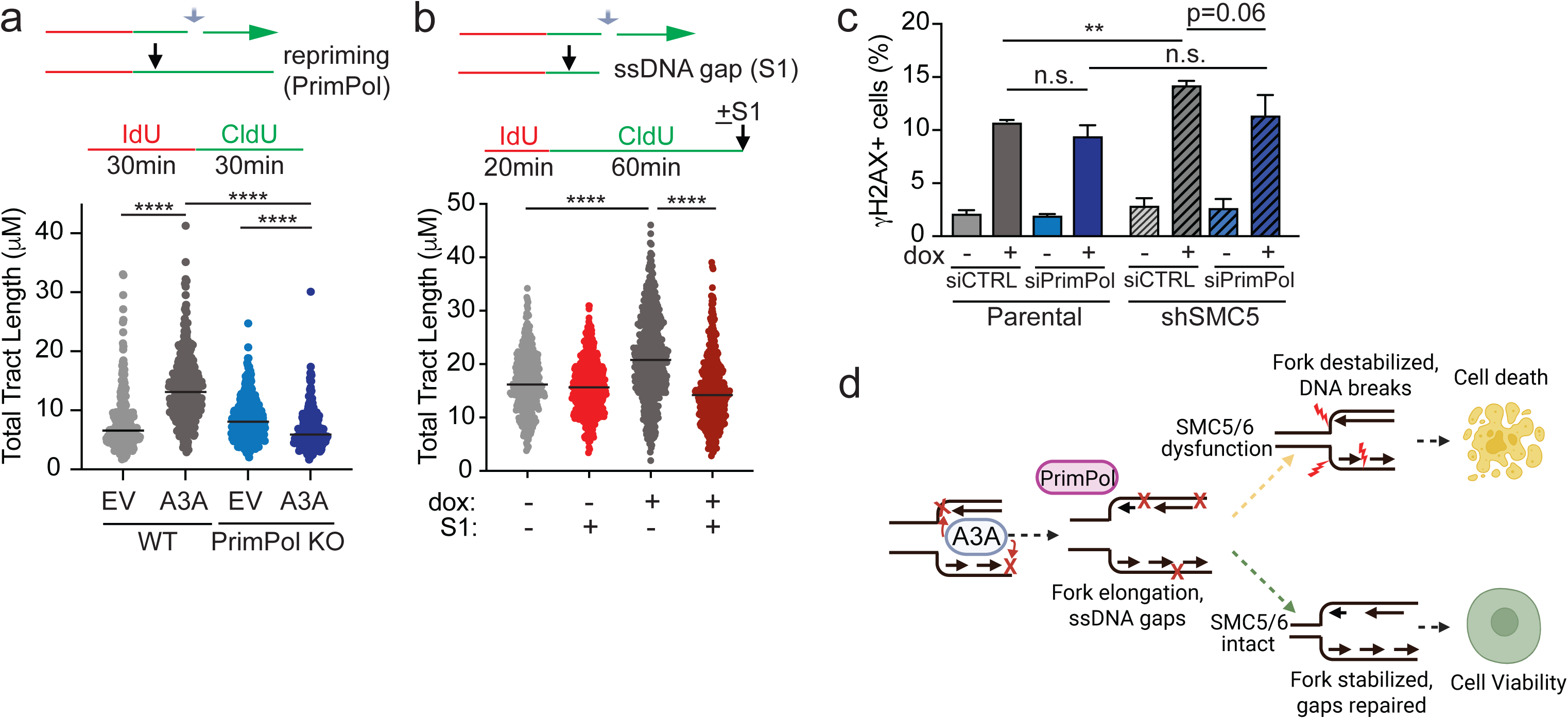
APOBEC3A-mediated replication fork elongation is dependent on PrimPol. (**a**) Depiction of replication fork lesion bypassed by PrimPol-mediated repriming downstream of the lesion (gray arrow), resulting in longer DNA fiber. Schematic of fiber assay. U2OS cells depleted of PrimPol by CRISPR-Cas9 editing were compared to isogenic controls. Total tract length (CldU+IdU) of DNA fibers was measured 24 hours after transfection of empty vector (EV) or APOBEC3A (A3A). (**b**) Depiction of replication fork lesion with ssDNA gap (gray arrow) which would be expected to result in shorter DNA fiber upon cleavage of the ssDNA gap by S1 nuclease. Schematic of S1 fiber assay. Total tract length (CldU+IdU) of U2OS-A3A cells treated with dox for 72 hours. DNA fiber assays were performed in biological triplicate and analyzed by Kruskal-Wallis test. Bars are median, ****p<0.0001. (**c**) K562-A3A cells were depleted of SMC5 by integrated shRNA and/or PrimPol by siRNA transfection. Cells were treated with dox for 72 hours then analyzed for γH2AX by flow cytometry. Error bars are SD of n=3 biological replicates. p-value by two-tailed t-test, **p<0.01. (**d**) Proposed model of APOBEC3A-induced genotoxicity enabled by SMC5/6 dysfunction.

PrimPol-mediated repriming leaves short ssDNA gaps behind the replication fork where damaged DNA was skipped (**Fig 7b**)(Quinet et al. 2017; Taglialatela et al. 2021; Tirman et al. 2021). Post-replicative gaps are too small to be visualized at the resolution of DNA fiber imaging(Quinet et al. 2017), therefore replication tracts undergoing PrimPol-mediated repriming should appear longer despite containing ssDNA gaps. To determine whether APOBEC3A activity results in post-replicative ssDNA gap formation, we used a modified version of the DNA fiber protocol in which genomic DNA is treated with an ssDNA-specific S1 endonuclease after pulse labeling with IdU and CldU(Quinet et al. 2017). Shorter DNA fibers result upon S1 treatment if ssDNA gaps are present (**Fig 7b**). Treatment with S1 nuclease led to significantly decreased DNA fiber length in cells with active APOBEC3A (**Fig 7b**). Our findings suggest that APOBEC3A likely does not cause an increased rate of DNA synthesis but rather causes apparent elongation of replication tracts due to “skipping” of base lesions by PrimPol.

We queried whether PrimPol loss would cause substantial DNA damage in cells that express APOBEC3A, similar to the phenomenon noted with SMC5/6 loss. Instead we found that cells depleted of PrimPol did not have increased levels of γH2AX when APOBEC3A was expressed (**Fig 7c, SF5a-b**). These data suggest that PrimPol mediates replication tract lengthening but not genome stability upon APOBEC3A-induced DNA lesions. Additionally, we found that simultaneous depletion of both PrimPol and SMC5 slightly decreased the number of cells with γH2AX staining in APOBEC3A-expressing cells relative to those with selective SMC5 depletion. These results suggest that PrimPol and SMC5 coordinate a response to APOBEC3A-induced DNA lesions at replication forks (**Fig 7d**).

## DISCUSSION

Tumor genome sequencing has demonstrated that mutagenesis from APOBEC3A is widespread throughout human cancers(Petljak and Alexandrov 2016; Alexandrov et al. 2020), however the mechanisms that enable dysregulated activity of APOBEC3A in cancer remain elusive. Several genomic determinants that enhance APOBEC3A activity have been elucidated recently, including a preference for acting at T<u>C</u> dinucleotides, stem-loop structures, and ssDNA at replication forks(Nik-Zainal et al. 2012; Seplyarskiy et al. 2016; Buisson et al. 2017; Buisson et al. 2019; Petljak et al. 2019; Jalili et al. 2020; Langenbucher et al. 2021). The mechanisms by which cells respond to APOBEC3A activity in order to maintain genome integrity have also been examined and include the replication checkpoint(Buisson et al. 2017; Green et al. 2017) as well as HMCES which protects abasic sites in ssDNA(Mehta et al. 2020; Biayna et al. 2021). These prior studies demonstrate that multiple genome-protective responses are required to prevent cytotoxicity from APOBEC3A. We now report a previously unknown, conserved mechanism of genome protection from APOBEC3A activity enacted by the SMC5/6 complex.

In addition to defining a dependence on SMC5/6, we found that cells with APOBEC3A expression exhibit elongated replication tracts relative to controls. This is counterintuitive to what would be expected of a response to base damage in ssDNA. Prior studies have found that conditions in which replication forks accelerate, such as PARP inhibition, are associated with genome instability and DNA damage(Zhong et al. 2013; Maya-Mendoza et al. 2018; Merchut-Maya et al. 2019). In contrast, lengthening of nascent strands in DNA fibers after APOBEC3A activity is likely due to extension of DNA synthesis beyond obstacles rather than an increased rate of DNA synthesis. While we found that longer replication tracts were generated after 24-72 hours of APOBEC3A expression, a prior study found that brief APOBEC3A expression within 4 hours led to decreased tract lengths(Mehta et al. 2020). We posit that the differences between these two findings reflect early versus late responses to APOBEC3A activity; deamination-induced damage may initially result in fork slowing or stalling, however the same forks may be able to progress upon activation or recruitment of DNA damage tolerance pathways. In fact, we found that fork lengthening in response to APOBEC3A was dependent on PrimPol, which may not be recruited to replication forks immediately upon deamination. Indeed, in a study of cisplatin-induced replication stress, PrimPol was found to be upregulated and chromatin-associated only upon treatment with a second dose of cisplatin(Quinet et al. 2020). We envision a similar time-dependent adaptation to APOBEC3A activity in PrimPol upregulation and recruitment to stalled forks with subsequent re-initiation of DNA synthesis.

The SMC5/6 complex has structural and catalytic functions, all of which have been demonstrated to play roles in genome stability(Aragon 2018; Peng and Zhao 2023). Although the function of SMC5/6 in genome maintenance is not fully understood, recent single-molecule studies demonstrate stable binding of Smc5/6 to ssDNA-dsDNA junctions, which mimic DNA replication and repair structures(Chang et al. 2022; Tanasie et al. 2022). This observation suggests that the DNA binding activity of Smc5/6 can be important for protecting junction-containing structures, which increase due to APOBEC3A activity during replication. Our yeast data support this idea. Thus, a potential model to explain the synthetic lethal interaction between APOBEC3A activity and loss of SMC5/6 is that deaminase activity at the replication fork leads to ssDNA gaps generated by PrimPol, which are protected from cleavage by SMC5/6 binding DNA. In this model, SMC5/6 loss can destabilize forks and gaps leading to DNA breaks and genotoxicity (**Fig 7d**). One mechanism by which SMC5/6 may protect ssDNA gaps is through homology-directed repair (HDR). Post-replicative gaps are repaired in part through template switching, a replication-specific HDR pathway(Tirman et al. 2021). SMC5/6 regulates resolution of HDR intermediates during replication-associated repair(Ampatzidou et al. 2006; Chen et al. 2009; Irmisch et al. 2009). Additionally, PrimPol has been proposed to prevent mutagenesis from APOBEC3 enzymes by stimulating HDR to limit error-prone DNA synthesis(Pilzecker et al. 2016). Thus, SMC5/6 may be important for a PrimPol-initiated HDR pathway.

This model is supported by several of our findings regarding coordination of genome protection by PrimPol and SMC5/6 upon APOBEC3A activity. First, we found that PrimPol loss alone in cells with active APOBEC3A does not cause increased DNA damage. Loss of Primpol would be predicted to decrease ssDNA gap generation and therefore mitigate the potential for cleavage leading to DNA breaks. Additionally, our data demonstrate that PrimPol loss partially rescues DNA damage caused by SMC5/6 loss. In the absence of PrimPol, fewer ssDNA gaps would be expected thus the need for SMC5/6 to prevent cleavage is diminished. Alternative mechanisms of ssDNA gap protection by SMC5/6, such as shielding from nucleases or stabilizing to prevent breaks, would also fit in this model.

A non-epistatic model is also possible in which PrimPol and SMC5/6 may promote fork protection independent of one another. PrimPol may not be the only DNA damage tolerance mechanism that is enabled by SMC5/6. Indeed, we find that yeast, which lack a PrimPol homologue, can tolerate expression of APOBEC3A as long as SMC5/6 is intact. In yeast, and perhaps also in mammalian cells, SMC5/6 may enable additional mechanisms of fork protection through fork reversal(Thompson and Cortez 2020) or gap filling(Peng and Feng 2016) as previously proposed. In the future, modeling of functional SMC5/6 mutants in mammalian cells may provide an opportunity to mechanistically examine consequences of SMC5/6 dysfunction.

While SMC5/6 has multiple functions, complex formation relies on all subunits being intact(Potts and Yu 2005; Gallego-Paez et al. 2014; Venegas et al. 2020). Germline defects in SMC5/6 subunits associated with human diseases provide insight into how compromise of a single SMC5/6 subunit disrupts genome stability. For example, SMC5/6-destabilizing mutations in NSMCE3 cause lung disease-immunodeficiency-chromosomal breakage syndrome (LICS)(van der Crabben et al. 2016), and NSMCE2 or SMC5 mutations result in primordial dwarfism and insulin resistance(Payne et al. 2014; Zhu et al. 2023). In this study, we find that deleterious mutations in SMC5/6 subunits correlate with high tumor mutational burdens in human cancers. Our findings are consistent with a recent *in silico* study showing that tumors with alterations in SMC5/6 genes display markers of genome instability, such as aneuploidy(Roy et al. 2023). These data raise questions regarding the etiology of mutagenesis and the sensitivity of those tumors to genotoxic agents. Interestingly, we find that SBS10 and SBS14, signatures consistent with DNA polymerase epsilon (pol ε) dysfunction, are overrepresented in SMC5/6-mutant tumors. Experimental data in yeast suggest that SUMOylation of pol ε by SMC5/6 promotes DNA synthesis(Meng et al. 2019; Winczura et al. 2019). Our findings in human cancers indicate a similar dependence of pol ε function on SMC5/6. Identification of additional mutational processes in SMC5/6-mutant tumors may provide insight into the contexts in which SMC5/6 dysfunction is permissive of mutagenesis. These studies may indicate opportunities to exploit mutagenesis in tumors with dysfunctional SMC5/6 as a therapeutic vulnerability.

## MATERIALS AND METHODS

### Cell culture and plasmid transfection

HCT116-NSE4A/SMC6-mAID, a kind gift from the lab of Ian Hickson(Venegas et al. 2020), U2OS-A3A, U2OS PrimPol knockout(Quinet et al. 2020), and 293T cells (used for lentiviral production) were maintained in DMEM media supplemented with 10% tetracycline-free FBS and 1% Pen-Strep. THP1, K562, and Jurkat cells were maintained in RPMI media supplemented with 10% tetracycline-free FBS and 1% Pen-Strep. All cells were grown at 37°C in a humidified atmosphere containing 5% CO2. Expression vectors containing APOBEC3A (pcDNA-A3A-GFP)(Landry et al. 2011) or GFP alone (pcDNA-GFP) were transfected using Lipofectamine 2000 (Thermo Fisher)

### Lentivectors and cell line generation

THP1-A3A, U2OS-A3A, and U2OS-C106S cells were generated by lentiviral transduction as previously described(Everett et al. 2009; Landry et al. 2011; Green et al. 2016; Green et al. 2017). Cas9 was introduced to THP1-A3A using lentivirus (lenti-Cas9-blast)(Sanjana et al. 2014). Blasticidin (Santa Cruz) selection began 24 hours after transduction until non-transduced controls were 100% non-viable. Inducible HCT116-NSE4A/SMC6-mAID-A3A were generated by lentiviral transduction using a dox-inducible pFLRU-A3A lentivector with Thy1.2 selection marker. Cells were bead sorted using magnetic anti-Thy1.2 beads (Miltenyi) until a stable >95% Thy1.2+ population was achieved. K562-A3A, K562-A3A*C106S, and Jurkat-A3A were generated by lentiviral transduction using the dox-inducible pSLIK-A3A lentivector with G418 resistance as previously described(Green et al. 2017). All A3A transgenes have a C-terminal HA tag.

### Genome-wide CRISPR-Cas9 knockout screen in THP1-A3A cells

#### Functional screen

Pooled lentivirus encoding the Brunello guide RNA library was generated as previously described(Shalem et al. 2014). Large-scale spinfection was carried out with the same conditions described above, using 12-well plates with 2×10^6^ cells per well. Each well was transduced with 50 μl Brunello library lentivirus. Wells were pooled into 15 cm plates after spinfection and overnight incubation and selected using puromycin for 7 days. Following puromycin selection, THP1-A3A-Cas9 cells were plated in triplicate into a 12-well plate at a concentration of 2×10^6^ cells per well. Doxycycline was added to +dox wells every 48 hours beginning on day 0. 400 x10^6^ cells cultured in parallel received vehicle control (water) at equal volume. Cells from dox-treated and untreated wells were harvested on the day of dox induction and after 15 days of dox treatment. Genomic DNA was extracted using Genomic DNA mini kit (Invitrogen) on a pre-PCR bench under sterile conditions to avoid DNA contamination.

#### Amplification and sequencing of library gRNAs

Guide RNAs were amplified by PCR from cellular genomic DNA and amplified using one-step PCR with barcodes on reverse primers, as previously described(Shalem et al. 2014). Illumina next-generation sequencing was applied to an amplicon generated from each integrated gRNA(Shalem et al. 2014). Briefly, we used all collected gDNA (1,000× coverage) divided into 100 μL PCR reactions with 5 μg of DNA per reaction. Takara ExTaq DNA Polymerase was used with the following PCR program: [95° 2 minutes (98° 10 seconds, 60° 30 seconds, 72° 30 seconds) × 24, 72° 5 minutes]. PCR products were gel-purified using the QiaQuick Gel Extraction Kit (Qiagen). Quality assessment was done by qubit (for concentration), bioAnalyzer (for size distribution), and Kapa Library Quantification (for clusterable molarity). The purified pooled library was then sequenced on a HiSeq4000 with ∼5% PhiX added to the sequencing lane.

#### Genome-Wide Screen Analysis

To count the number of reads associated with each sgRNA taken from the raw Fastq file, we first extracted the sgRNA targeting sequencing using a regular expression containing the three nucleotides flanking each side of the sgRNA 20 bp target. sgRNA spacer sequences were then aligned to a preindexed Brunello library (Addgene) using the short-read aligner “bowtie” using parameters (-v 0 -m 1). Data analysis was performed using custom R scripts. Table 1 shows the output only for MAGeCK version 0.5.9.5 where the counts were normalized to the 1000 non-targeting control sgRNA contained in the Brunello library.

### Gene depletion by RNAi

#### Short-hairpin RNA

Commercially available shRNA lentiviral vectors (Sigma, TRCN0000147948, TRCN0000148162, TRCN0000147348, TRCN0000147918) were used to construct SMC5-depleted K562-A3A and Jurkat-A3A cell lines. Cells were infected with shRNA lentivirus, selected in 1 mg/mL puromycin until non-transduced controls were 100% non-viable. Of note, constitutive shRNA-mediated depletion of SMC5 was lost after 3-4 months in culture. Many vials were frozen after each cell line generation to preserve those with maximal gene depletion. Cells were only cultured for use in experiments for <6 weeks. Confirmation of gene depletion was done at least monthly while cells were in culture.

#### Small interfering RNA

Pooled siRNA oligonucleotides (25pmol) targeting SMC5 (Horizon SMARTpool) were transfected into 1×10^6^ cells using the RNAiMAX transfection reagent (Invitrogen) according to the manufacturer’s protocol. Gene depletion was confirmed by immunoblot and/or quantitative PCR.

### Antibodies

Commercially available antibodies used for immunoblotting, immunofluorescence, intracellular staining, and DNA fiber spreading were obtained from Santa Cruz Biotechnology (Tubulin, Ku86, SMC5, SMC6), GeneTex (SMC5), Novus Biologicals (Cas9 Antibody 7A9-3A3), Abcam (RPA, BrdU), Biolegend (HA), Cell Signaling (HA, γH2AX, cyclin A, pChk1 S317), Invitrogen (PGK1), the NIH AIDS Reagent Program (APOBEC3A/B), and BD Biosciences (γH2AX-488, γH2AX-647, BrdU). Secondary antibodies for immunoblotting were obtained from Jackson ImmunoResearch (goat anti-rabbit IgG, goat anti-mouse IgG). Secondary antibodies for immunofluorescence were obtained from Invitrogen (Alexa Fluor 488 goat anti-mouse IgG, Alexa Fluor 568 goat anti-rabbit IgG). Secondary antibodies for DNA fiber spreading were obtained from Invitrogen (Alexa Fluor 488 chicken anti-rat IgG, Alexa Fluor 546 goat anti-mouse IgG).

### Viability assays

To assess proportions of live and dead cells, staining was performed using the Live/Dead Kit (Invitrogen) according to the manufacturer’s instructions. Data were collected using a Fortessa Flow Cytometer (BD Biosciences) or Accuri C6 Flow Cytometer (BD Biosciences) and analyzed by FlowJo software. To assess viability by metabolic activity, cells were plated in triplicate in a 96-well plate, precultured for 24 hours, then 50 μl media with and without doxycycline was added to each well every other day. 10 μl WST-8 reagent from the Cell Counting Kit-8 (Dojindo) was added to each well 4-6 hours prior to analysis using a microplate reader (BMG Labtech Omega).

### Proliferation Assay

On day 0, cells were plated at a density of 200,000 cells per well in a 6-well plate. Each cell type was grown in the presence and absence of 1 μg/mL doxycycline. Cells grown in the presence of doxycycline received doxycycline doses every other day. On days 3, 5, and 7, data were collected using an automatic cell counter (Countess, ThermoFisher).

### Intracellular γH2AX detection by flow cytometry

Cells were harvested, fixed and permeabilized using reagents from the CytoFix/CytoPerm Kit (BD Biosciences) according to the manufacturer’s instructions. Cells were stained with a fluorophore-conjugated γH2AX antibody (BD AlexaFluor 488 or 647 Mouse Anti-H2AX (pS139)) at a ratio of 10 μl antibody per 100 μl cells (<1×10^6^ cells/sample). Data were collected using a Fortessa Flow Cytometer (BD Biosciences) or Accuri C6 Flow Cytometer (BD Biosciences) and analyzed by FlowJo software.

### Cell Synchronization and Cell Cycle Analysis

Cell synchronization was achieved by double thymidine block as previously described(Chen and Deng 2018), with the following minor modifications: 2mM thymidine was added to cells for 24 hours then removed by change of media. After 9 hours recovery, thymidine was again added for 24 hours. Following removal of second thymidine pulse, cells were analyzed at time 0 and released into thymidine-free media. To analyze cell cycle, cells were fixed in 70% ice cold ethanol, washed in PBS, and resuspended in staining solution containing Triton X, RNAseA, and 1 mg/mL propidium iodide (Biotum). Data were collected using an Accuri C6 Flow Cytometer and analyzed by FlowJo software.

### Immunoblotting and Immunofluorescence

Cell lysates were prepared by harvesting cells in LDS buffer and boiling for 15 minutes, then adding 20% β−mercaptoethanol. Lysates were run on Bis-Tris gels and transferred to a nitrocellulose membrane. After incubation with primary and secondary antibodies, membranes were developed using ECL Western blotting reagents (Pierce) on a GelDoc Go system (BioRad). For immunofluorescence, cells were cultured on cover slips. Following treatment, cells were pre-extracted using 0.5% Triton-X in PBS for 15 minutes on ice to visualize chromatin-bound proteins (i.e. RPA). All other immunofluorescence experiments proceeded as followed: cells were fixed with 4% paraformaldehyde for 15 minutes at room temperature (RT), permeabilized with 0.5% Triton-X for 10 minutes at RT, and blocked with 5% BSA for 1 hour at RT. Primary antibodies were diluted in 5% BSA and incubated with slides for 1 hour to overnight. Secondary antibodies used were anti-mouse or rabbit Alexa Fluor 488 and 568 (BD Biosciences). Nuclei were visualized by 4.6-diamidino-2-phenylindole (DAPI, ThermoFisher). Images were acquired using an inverted fluorescent microscope with attached camera (Leica) and processed using ImageJ. Protein foci and cell staining were analyzed in a blinded fashion.

### Comet assay

Neutral comet assays were performed using CometAssay (Trevigen) according to the manufacturer’s protocol with minor modifications as previously described(Wood et al. 2020). In brief, cells were harvested and resuspended at 3×10^5^ cells/mL in ice cold PBS, combined with molten LMAgarose, plated onto a comet slide, and allowed to dry at 4°C. Slides were incubated in lysis solution for 1 hour at 4°C and then immersed in 1X TBE buffer for 30 minutes at 4°C. Then slides underwent electrophoresis at 25 V for 30-45 minutes at 4°C in 1X TBE buffer. After electrophoresis, slides were washed in DNA precipitation solution (1 M ammonium acetate, 95% ethanol) and fixed in 70% ethanol for 30 minutes. Fixed slides were dried overnight at room temperature in the dark, stained with 1X SYBR Gold (Applied Biosystems), and washed twice with water. Images were acquired using a fluorescence microscope (Leica). Images were scored using the OpenComet plugin in ImageJ.

### Yeast strains and genetic techniques

All strains used are in W303 background *(ade2-1 can1-100 ura3-1 his3-11,15, leu2-3, 112 trp1-1 rad5-535)* containing wild-type *RAD5*. *smc5-DNAm* and *nse4-DNAm* strains are from Yu, et al(Yu et al. 2022). APOBCE3A expression plasmid containing hygromycin drug resistant marker is a derivative of pySR419-A3A(Hoopes et al. 2016) (gifted from Dr. Steven A. Roberts). This plasmid and its control vector were transformed into yeast cells individually using standard method and cells were grown on YPD plates containing hygromycin (300 μg/mL) at 30°C for 48 hours. Cell growth for three independent transformants in each case were assessed at 30°C and 37°C.

### DNA Fiber assay

U2OS and HCT116 cells were first pulse labeled for 30 minutes with 20μM IdU, washed three times with 1X DPBS, and then pulsed with 100 μM CldU for 30 minutes. For K562 and Jurkat cells, cells were first pulsed with 20μM IdU and then flushed with 100μM CldU for 30 minutes. After pulse, cells were harvested and collected in ice cold DPBS (∼1500 cells/μL). For the DNA fiber assay with the ssDNA-specific S1 nuclease (S1 Fiber), cells were permeabilized with CSK100 (100 mM NaCl, 10 mM MOPS pH 7, 3 mM MgCl_2_, 300 mM sucrose and 0.5% Triton X-100 in water) after the CldU pulse for 10 minutes at room temperature, treated with the S1 nuclease (ThermoFisher Scientific) at 20 U/mL in S1 buffer (30 mM sodium acetate pH 4.6, 10 mM zinc acetate, 5% glycerol, 50 mM NaCl in water) for 30 minutes at 37°C, and collected in PBS-0.1%BSA with cell scraper. Nuclei were then pelleted at ∼4600 x g for 5 minutes at 4°C, then resuspended in PBS (nuclei cannot be quantified, so initial number of cells plated should be considered when resuspending to a final concentration of 1,500 nuclei/μl). To spread fibers, 2 μL of cell solution was placed on charged glass slide, mixed with 6 μL of lysis buffer (200 mM Tris-HCl pH 7.4, 0.5% SDS, 50 mM EDTA), and gravity was used to spread DNA fibers. DNA fibers were fixed in a 3:1 solution of methanol and acetic acid, denatured in 2.5M HCl for 1 hour, and blocked in pre-warmed 5% BSA at 37°C for 1 hour. IdU and CldU were detected using mouse anti-BrdU (1:20, Invitrogen) and rat anti-BrdU (1:75, Abcam) respectively for 1.5 hours at room temperature in a humid chamber followed by anti-mouse Alexa-546 (1:50) and anti-rat Alexa-488 (1:50) for 1 hour at room temperature in a humid chamber. Slides were mounted with Prolong Gold Antifade Solution (Invitrogen) and cured overnight at room temperature protected from light. Fibers were imaged with a 63X oil objective on a Leica DM4 B. Quantification and measurement of fibers was done in Image J by blinded analysis.

### Mutational signature analysis

Mutation calls of the SMC5/6 complex genes were obtained from the Genomic Data Commons Data Portal at https://docs.gdc.cancer.gov (v.36)(Grossman et al. 2016). Those predicted to cause negative effects on the proteins’ functions by either Ensembl VEP(McLaren et al. 2016) or SIFT(Choi et al. 2012) were classified as deleterious mutations. Additionally, single base substitutions (SBSs) of 9493 TCGA tumors were obtained from the COSMIC database at https://www.synapse.org/#!Synapse:syn11726601/files. Tumors with one or more deleterious mutations in SMC5/6 complex were defined by merging these two datasets and were later used in downstream analysis. In contrast, other genes mutated in tumors with intact SMC5/6 complex were used as a comparison gene set (**Supplemental Table 2**). A set of control tumors was then defined as those which carried detrimental mutations of the comparison gene set. Differences in mutational burden and APOBEC enrichment between samples were inspected and visualized using the R packages ggplot2 and tidyverse, while statistical difference was accessed by two-sided Mann-Whitney test. The R package MutationalPatterns(Manders et al. 2022) was used to study the contribution of other COSMIC SBS signatures (v3.2).

### Statistical analysis

All statistical tests were performed in R or GraphPad (Prism). Biological and/or technical triplicate tests were used to ensure robustness and reproducibility of data. Standard deviations, standard error of the mean, and p values were generated using paired and unpaired two-tail t-tests, F-tests, or Anova.

### Data availability

All data presented in this manuscript are available from the corresponding author upon request.

## ACKNOWLEDGEMENTS

We thank Dr. Jieya Shao, Dr. Sebastien Landry, and all members of the Green Lab for helpful discussions and input. We thank Stephen Sykes, Lianna Valin, Ian Hickson, Luis Batista, and members of the Vindigni and Bednarski Labs for sharing reagents and technical support. Special thanks to Dr. Steven Roberts for providing expression plasmids and advice regarding APOBEC3A expression in yeast. This study was funded by support from the NIH T32 HL007088 (DFF), NIH K08 CA212299 (AMG), DOD CA200867 (AMG), Cancer Research Foundation, Children’s Discovery Institute, and American Cancer Society. Work in the AV lab was supported by the National Cancer Institute (NCI) grants R01CA237263 and R01CA248526.

## AUTHOR CONTRIBUTIONS

DRO, ARH, DFF, BRH, MT, RAD, AM, JHS, JD, JF, and EC performed experiments, analyzed and interpreted data. TT and KEH performed informatic experiments and statistical tests. DRO, ARH, DFF, AMG, MDW, OS, JB, XZ and AV conceptualized and designed experiments. AMG wrote the manuscript, with editing from all authors.

## REFERENCES

Agashe S, Joseph CR, Reyes TAC, Menolfi D, Giannattasio M, Waizenegger A, Szakal B, Branzei D. 2021. Smc5/6 functions with Sgs1-Top3-Rmi1 to complete chromosome replication at natural pause sites. Nat Commun 12: 2111.

Alabert C, Bukowski-Wills JC, Lee SB, Kustatscher G, Nakamura K, de Lima Alves F, Menard P, Mejlvang J, Rappsilber J, Groth A. 2014. Nascent chromatin capture proteomics determines chromatin dynamics during DNA replication and identifies unknown fork components. Nat Cell Biol 16: 281–293.

Alexandrov LB, Kim J, Haradhvala NJ, Huang MN, Tian Ng AW, Wu Y, Boot A, Covington KR, Gordenin DA, Bergstrom EN et al. 2020. The repertoire of mutational signatures in human cancer. Nature 578: 94–101.

Alt A, Dang HQ, Wells OS, Polo LM, Smith MA, McGregor GA, Welte T, Lehmann AR, Pearl LH, Murray JM et al. 2017. Specialized interfaces of Smc5/6 control hinge stability and DNA association. Nat Commun 8: 14011.

Ampatzidou E, Irmisch A, O’Connell MJ, Murray JM. 2006. Smc5/6 is required for repair at collapsed replication forks. Mol Cell Biol 26: 9387–9401.

Aragon L. 2018. The Smc5/6 Complex: New and Old Functions of the Enigmatic Long-Distance Relative. Annu Rev Genet 52: 89–107.

Atkins A, Xu MJ, Li M, Rogers NP, Pryzhkova MV, Jordan PW. 2020. SMC5/6 is required for replication fork stability and faithful chromosome segregation during neurogenesis. Elife 9.

Baker SC, Mason AS, Slip RG, Skinner KT, Macdonald A, Masood O, Harris RS, Fenton TR, Periyasamy M, Ali S et al. 2022. Induction of APOBEC3-mediated genomic damage in urothelium implicates BK polyomavirus (BKPyV) as a hit-and-run driver for bladder cancer. Oncogene 41: 2139–2151.

Barlow JH, Faryabi RB, Callen E, Wong N, Malhowski A, Chen HT, Gutierrez-Cruz G, Sun HW, McKinnon P, Wright G et al. 2013. Identification of early replicating fragile sites that contribute to genome instability. Cell 152: 620–632.

Betts Lindroos H, Strom L, Itoh T, Katou Y, Shirahige K, Sjogren C. 2006. Chromosomal association of the Smc5/6 complex reveals that it functions in differently regulated pathways. Mol Cell 22: 755–767.

Biayna J, Garcia-Cao I, Alvarez MM, Salvadores M, Espinosa-Carrasco J, McCullough M, Supek F, Stracker TH. 2021. Loss of the abasic site sensor HMCES is synthetic lethal with the activity of the APOBEC3A cytosine deaminase in cancer cells. PLoS Biol 19: e3001176.

Buisson R, Langenbucher A, Bowen D, Kwan EE, Benes CH, Zou L, Lawrence MS. 2019. Passenger hotspot mutations in cancer driven by APOBEC3A and mesoscale genomic features. Science 364.

Buisson R, Lawrence MS, Benes CH, Zou L. 2017. APOBEC3A and APOBEC3B Activities Render Cancer Cells Susceptible to ATR Inhibition. Cancer Res 77: 4567–4578.

Burns MB, Lackey L, Carpenter MA, Rathore A, Land AM, Leonard B, Refsland EW, Kotandeniya D, Tretyakova N, Nikas JB et al. 2013a. APOBEC3B is an enzymatic source of mutation in breast cancer. Nature 494: 366–370.

Burns MB, Temiz NA, Harris RS. 2013b. Evidence for APOBEC3B mutagenesis in multiple human cancers. Nat Genet 45: 977–983.

Chan K, Roberts SA, Klimczak LJ, Sterling JF, Saini N, Malc EP, Kim J, Kwiatkowski DJ, Fargo DC, Mieczkowski PA et al. 2015. An APOBEC3A hypermutation signature is distinguishable from the signature of background mutagenesis by APOBEC3B in human cancers. Nat Genet 47: 1067–1072.

Chang JT, Li S, Beckwitt EC, Than T, Haluska C, Chandanani J, O’Donnell ME, Zhao X, Liu S. 2022. Smc5/6’s multifaceted DNA binding capacities stabilize branched DNA structures. Nat Commun 13: 7179.

Chen G, Deng X. 2018. Cell Synchronization by Double Thymidine Block. Bio Protoc 8.

Chen H, Lilley CE, Yu Q, Lee DV, Chou J, Narvaiza I, Landau NR, Weitzman MD. 2006. APOBEC3A is a potent inhibitor of adeno-associated virus and retrotransposons. Curr Biol 16: 480–485.

Chen YH, Choi K, Szakal B, Arenz J, Duan X, Ye H, Branzei D, Zhao X. 2009. Interplay between the Smc5/6 complex and the Mph1 helicase in recombinational repair. Proc Natl Acad Sci U S A 106: 21252–21257.

Choi Y, Sims GE, Murphy S, Miller JR, Chan AP. 2012. Predicting the functional effect of amino acid substitutions and indels. PLoS One 7: e46688.

Cortez LM, Brown AL, Dennis MA, Collins CD, Brown AJ, Mitchell D, Mertz TM, Roberts SA. 2019. APOBEC3A is a prominent cytidine deaminase in breast cancer. PLoS Genet 15: e1008545.

DeWeerd RA, Nemeth E, Poti A, Petryk N, Chen CL, Hyrien O, Szuts D, Green AM. 2022. Prospectively defined patterns of APOBEC3A mutagenesis are prevalent in human cancers. Cell Rep 38: 110555.

Doench JG, Fusi N, Sullender M, Hegde M, Vaimberg EW, Donovan KF, Smith I, Tothova Z, Wilen C, Orchard R et al. 2016. Optimized sgRNA design to maximize activity and minimize off-target effects of CRISPR-Cas9. Nat Biotechnol 34: 184–191.

Elango R, Osia B, Harcy V, Malc E, Mieczkowski PA, Roberts SA, Malkova A. 2019. Repair of base damage within break-induced replication intermediates promotes kataegis associated with chromosome rearrangements. Nucleic Acids Res 47: 9666–9684.

Everett RD, Parsy ML, Orr A. 2009. Analysis of the functions of herpes simplex virus type 1 regulatory protein ICP0 that are critical for lytic infection and derepression of quiescent viral genomes. J Virol 83: 4963–4977.

Gallego-Paez LM, Tanaka H, Bando M, Takahashi M, Nozaki N, Nakato R, Shirahige K, Hirota T. 2014. Smc5/6-mediated regulation of replication progression contributes to chromosome assembly during mitosis in human cells. Mol Biol Cell 25: 302–317.

Grange LJ, Reynolds JJ, Ullah F, Isidor B, Shearer RF, Latypova X, Baxley RM, Oliver AW, Ganesh A, Cooke SL et al. 2022. Pathogenic variants in SLF2 and SMC5 cause segmented chromosomes and mosaic variegated hyperploidy. Nat Commun 13: 6664.

Green AM, Budagyan K, Hayer KE, Reed MA, Savani MR, Wertheim GB, Weitzman MD. 2017. Cytosine Deaminase APOBEC3A Sensitizes Leukemia Cells to Inhibition of the DNA Replication Checkpoint. Cancer Res 77: 4579–4588.

Green AM, Landry S, Budagyan K, Avgousti DC, Shalhout S, Bhagwat AS, Weitzman MD. 2016. APOBEC3A damages the cellular genome during DNA replication. Cell Cycle 15: 998–1008.

Grossman RL, Heath AP, Ferretti V, Varmus HE, Lowy DR, Kibbe WA, Staudt LM. 2016. Toward a Shared Vision for Cancer Genomic Data. N Engl J Med 375: 1109–1112.

Haradhvala NJ, Polak P, Stojanov P, Covington KR, Shinbrot E, Hess JM, Rheinbay E, Kim J, Maruvka YE, Braunstein LZ et al. 2016. Mutational Strand Asymmetries in Cancer Genomes Reveal Mechanisms of DNA Damage and Repair. Cell 164: 538–549.

Harris RS, Dudley JP. 2015. APOBECs and virus restriction. Virology 479-480: 131-145.

Hoopes JI, Cortez LM, Mertz TM, Malc EP, Mieczkowski PA, Roberts SA. 2016. APOBEC3A and APOBEC3B Preferentially Deaminate the Lagging Strand Template during DNA Replication. Cell Rep 14: 1273–1282.

Irmisch A, Ampatzidou E, Mizuno K, O’Connell MJ, Murray JM. 2009. Smc5/6 maintains stalled replication forks in a recombination-competent conformation. EMBO J 28: 144–155.

Jalili P, Bowen D, Langenbucher A, Park S, Aguirre K, Corcoran RB, Fleischman AG, Lawrence MS, Zou L, Buisson R. 2020. Quantification of ongoing APOBEC3A activity in tumor cells by monitoring RNA editing at hotspots. Nat Commun 11: 2971.

Ju L, Wing J, Taylor E, Brandt R, Slijepcevic P, Horsch M, Rathkolb B, Racz I, Becker L, Hans W et al. 2013. SMC6 is an essential gene in mice, but a hypomorphic mutant in the ATPase domain has a mild phenotype with a range of subtle abnormalities. DNA Repair (Amst) 12: 356–366.

Khan S, Ahamad N, Bhadra S, Xu Z, Xu YJ. 2022. Smc5/6 Complex Promotes Rad3(ATR) Checkpoint Signaling at the Perturbed Replication Fork through Sumoylation of the RecQ Helicase Rqh1. Mol Cell Biol 42: e0004522.

Landry S, Narvaiza I, Linfesty DC, Weitzman MD. 2011. APOBEC3A can activate the DNA damage response and cause cell-cycle arrest. EMBO Rep 12: 444–450.

Langenbucher A, Bowen D, Sakhtemani R, Bournique E, Wise JF, Zou L, Bhagwat AS, Buisson R, Lawrence MS. 2021. An extended APOBEC3A mutation signature in cancer. Nat Commun 12: 1602.

Law EK, Levin-Klein R, Jarvis MC, Kim H, Argyris PP, Carpenter MA, Starrett GJ, Temiz NA, Larson LK, Durfee C et al. 2020. APOBEC3A catalyzes mutation and drives carcinogenesis in vivo. J Exp Med 217.

Lehmann AR. 2005. The role of SMC proteins in the responses to DNA damage. DNA Repair (Amst) 4: 309–314.

Manders F, Brandsma AM, de Kanter J, Verheul M, Oka R, van Roosmalen MJ, van der Roest B, van Hoeck A, Cuppen E, van Boxtel R. 2022. MutationalPatterns: the one stop shop for the analysis of mutational processes. BMC Genomics 23: 134.

Maya-Mendoza A, Moudry P, Merchut-Maya JM, Lee M, Strauss R, Bartek J. 2018. High speed of fork progression induces DNA replication stress and genomic instability. Nature 559: 279–284.

McLaren W, Gil L, Hunt SE, Riat HS, Ritchie GR, Thormann A, Flicek P, Cunningham F. 2016. The Ensembl Variant Effect Predictor. Genome Biol 17: 122.

Mehta KPM, Lovejoy CA, Zhao R, Heintzman DR, Cortez D. 2020. HMCES Maintains Replication Fork Progression and Prevents Double-Strand Breaks in Response to APOBEC Deamination and Abasic Site Formation. Cell Rep 31: 107705.

Meng X, Wei L, Peng XP, Zhao X. 2019. Sumoylation of the DNA polymerase epsilon by the Smc5/6 complex contributes to DNA replication. PLoS Genet 15: e1008426.

Menolfi D, Delamarre A, Lengronne A, Pasero P, Branzei D. 2015. Essential Roles of the Smc5/6 Complex in Replication through Natural Pausing Sites and Endogenous DNA Damage Tolerance. Mol Cell 60: 835–846.

Merchut-Maya JM, Bartek J, Maya-Mendoza A. 2019. Regulation of replication fork speed: Mechanisms and impact on genomic stability. DNA Repair (Amst) 81: 102654.

Mouron S, Rodriguez-Acebes S, Martinez-Jimenez MI, Garcia-Gomez S, Chocron S, Blanco L, Mendez J. 2013. Repriming of DNA synthesis at stalled replication forks by human PrimPol. Nat Struct Mol Biol 20: 1383–1389.

Natsume T, Kiyomitsu T, Saga Y, Kanemaki MT. 2016. Rapid Protein Depletion in Human Cells by Auxin-Inducible Degron Tagging with Short Homology Donors. Cell Rep 15: 210–218.

Nik-Zainal S, Alexandrov LB, Wedge DC, Van Loo P, Greenman CD, Raine K, Jones D, Hinton J, Marshall J, Stebbings LA et al. 2012. Mutational processes molding the genomes of 21 breast cancers. Cell 149: 979–993.

Payne F, Colnaghi R, Rocha N, Seth A, Harris J, Carpenter G, Bottomley WE, Wheeler E, Wong S, Saudek V et al. 2014. Hypomorphism in human NSMCE2 linked to primordial dwarfism and insulin resistance. J Clin Invest 124: 4028–4038.

Peng J, Feng W. 2016. Incision of damaged DNA in the presence of an impaired Smc5/6 complex imperils genome stability. Nucleic Acids Res 44: 10216–10229.

Peng XP, Lim S, Li S, Marjavaara L, Chabes A, Zhao X. 2018. Acute Smc5/6 depletion reveals its primary role in rDNA replication by restraining recombination at fork pausing sites. PLoS Genet 14: e1007129.

Peng XP, Zhao X. 2023. The multi-functional Smc5/6 complex in genome protection and disease. Nat Struct Mol Biol 30: 724–734.

Petljak M, Alexandrov LB. 2016. Understanding mutagenesis through delineation of mutational signatures in human cancer. Carcinogenesis 37: 531–540.

Petljak M, Alexandrov LB, Brammeld JS, Price S, Wedge DC, Grossmann S, Dawson KJ, Ju YS, Iorio F, Tubio JMC et al. 2019. Characterizing Mutational Signatures in Human Cancer Cell Lines Reveals Episodic APOBEC Mutagenesis. Cell 176: 1282–1294 e1220.

Petljak M, Dananberg A, Chu K, Bergstrom EN, Striepen J, von Morgen P, Chen Y, Shah H, Sale JE, Alexandrov LB et al. 2022. Mechanisms of APOBEC3 mutagenesis in human cancer cells. Nature 607: 799–807.

Pilzecker B, Buoninfante OA, Pritchard C, Blomberg OS, Huijbers IJ, van den Berk PC, Jacobs H. 2016. PrimPol prevents APOBEC/AID family mediated DNA mutagenesis. Nucleic Acids Res 44: 4734–4744.

Potts PR, Yu H. 2005. Human MMS21/NSE2 is a SUMO ligase required for DNA repair. Mol Cell Biol 25: 7021–7032.

Quinet A, Carvajal-Maldonado D, Lemacon D, Vindigni A. 2017. DNA Fiber Analysis: Mind the Gap! Methods Enzymol 591: 55–82.

Quinet A, Tirman S, Cybulla E, Meroni A, Vindigni A. 2021. To skip or not to skip: choosing repriming to tolerate DNA damage. Mol Cell 81: 649–658.

Quinet A, Tirman S, Jackson J, Svikovic S, Lemacon D, Carvajal-Maldonado D, Gonzalez-Acosta D, Vessoni AT, Cybulla E, Wood M et al. 2020. PRIMPOL-Mediated Adaptive Response Suppresses Replication Fork Reversal in BRCA-Deficient Cells. Mol Cell 77: 461–474 e469.

Richardson SR, Narvaiza I, Planegger RA, Weitzman MD, Moran JV. 2014. APOBEC3A deaminates transiently exposed single-strand DNA during LINE-1 retrotransposition. Elife 3: e02008.

Roberts SA, Lawrence MS, Klimczak LJ, Grimm SA, Fargo D, Stojanov P, Kiezun A, Kryukov GV, Carter SL, Saksena G et al. 2013. An APOBEC cytidine deaminase mutagenesis pattern is widespread in human cancers. Nat Genet 45: 970–976.

Roberts SA, Sterling J, Thompson C, Harris S, Mav D, Shah R, Klimczak LJ, Kryukov GV, Malc E, Mieczkowski PA et al. 2012. Clustered mutations in yeast and in human cancers can arise from damaged long single-strand DNA regions. Mol Cell 46: 424–435.

Roy S, Zaker A, Mer A, D’Amours D. 2023. Large-scale phenogenomic analysis of human cancers uncovers frequent alterations affecting SMC5/6 complex components in breast cancer. NAR Cancer 5: zcad047.

Sanjana NE, Shalem O, Zhang F. 2014. Improved vectors and genome-wide libraries for CRISPR screening. Nat Methods 11: 783–784.

Seplyarskiy VB, Soldatov RA, Popadin KY, Antonarakis SE, Bazykin GA, Nikolaev SI. 2016. APOBEC-induced mutations in human cancers are strongly enriched on the lagging DNA strand during replication. Genome Res 26: 174–182.

Shalem O, Sanjana NE, Hartenian E, Shi X, Scott DA, Mikkelson T, Heckl D, Ebert BL, Root DE, Doench JG et al. 2014. Genome-scale CRISPR-Cas9 knockout screening in human cells. Science 343: 84–87.

Sobczak-Thepot J, Harper F, Florentin Y, Zindy F, Brechot C, Puvion E. 1993. Localization of cyclin A at the sites of cellular DNA replication. Exp Cell Res 206: 43–48.

Suspene R, Aynaud MM, Guetard D, Henry M, Eckhoff G, Marchio A, Pineau P, Dejean A, Vartanian JP, Wain-Hobson S. 2011. Somatic hypermutation of human mitochondrial and nuclear DNA by APOBEC3 cytidine deaminases, a pathway for DNA catabolism. Proc Natl Acad Sci U S A 108: 4858–4863.

Taglialatela A, Leuzzi G, Sannino V, Cuella-Martin R, Huang JW, Wu-Baer F, Baer R, Costanzo V, Ciccia A. 2021. REV1-Polzeta maintains the viability of homologous recombination-deficient cancer cells through mutagenic repair of PRIMPOL-dependent ssDNA gaps. Mol Cell 81: 4008–4025 e4007.

Tanasie NL, Gutierrez-Escribano P, Jaklin S, Aragon L, Stigler J. 2022. Stabilization of DNA fork junctions by Smc5/6 complexes revealed by single-molecule imaging. Cell Rep 41: 111778.

Taylor BJ, Nik-Zainal S, Wu YL, Stebbings LA, Raine K, Campbell PJ, Rada C, Stratton MR, Neuberger MS. 2013. DNA deaminases induce break-associated mutation showers with implication of APOBEC3B and 3A in breast cancer kataegis. Elife 2: e00534.

Thompson PS, Cortez D. 2020. New insights into abasic site repair and tolerance. DNA Repair (Amst) 90: 102866.

Tirman S, Quinet A, Wood M, Meroni A, Cybulla E, Jackson J, Pegoraro S, Simoneau A, Zou L, Vindigni A. 2021. Temporally distinct post-replicative repair mechanisms fill PRIMPOL-dependent ssDNA gaps in human cells. Mol Cell 81: 4026–4040 e4028.

Torres-Rosell J, Sunjevaric I, De Piccoli G, Sacher M, Eckert-Boulet N, Reid R, Jentsch S, Rothstein R, Aragon L, Lisby M. 2007. The Smc5-Smc6 complex and SUMO modification of Rad52 regulates recombinational repair at the ribosomal gene locus. Nat Cell Biol 9: 923–931.

van der Crabben SN, Hennus MP, McGregor GA, Ritter DI, Nagamani SC, Wells OS, Harakalova M, Chinn IK, Alt A, Vondrova L et al. 2016. Destabilized SMC5/6 complex leads to chromosome breakage syndrome with severe lung disease. J Clin Invest 126: 2881–2892.

Venegas AB, Natsume T, Kanemaki M, Hickson ID. 2020. Inducible Degradation of the Human SMC5/6 Complex Reveals an Essential Role Only during Interphase. Cell Rep 31: 107533.

Venkatesan S, Angelova M, Puttick C, Zhai H, Caswell DR, Lu WT, Dietzen M, Galanos P, Evangelou K, Bellelli R et al. 2021. Induction of APOBEC3 Exacerbates DNA Replication Stress and Chromosomal Instability in Early Breast and Lung Cancer Evolution. Cancer Discov 11: 2456–2473.

Wang L, Chen H, Wang C, Hu Z, Yan S. 2018. Negative regulator of E2F transcription factors links cell cycle checkpoint and DNA damage repair. Proc Natl Acad Sci U S A 115: E3837–E3845.

Winczura A, Appanah R, Tatham MH, Hay RT, De Piccoli G. 2019. The S phase checkpoint promotes the Smc5/6 complex dependent SUMOylation of Pol2, the catalytic subunit of DNA polymerase epsilon. PLoS Genet 15: e1008427.

Wood M, Quinet A, Lin YL, Davis AA, Pasero P, Ayala YM, Vindigni A. 2020. TDP-43 dysfunction results in R-loop accumulation and DNA replication defects. J Cell Sci 133.

Wu N, Kong X, Ji Z, Zeng W, Potts PR, Yokomori K, Yu H. 2012. Scc1 sumoylation by Mms21 promotes sister chromatid recombination through counteracting Wapl. Genes Dev 26: 1473–1485.

Yu Y, Li S, Ser Z, Kuang H, Than T, Guan D, Zhao X, Patel DJ. 2022. Cryo-EM structure of DNA-bound Smc5/6 reveals DNA clamping enabled by multi-subunit conformational changes. Proc Natl Acad Sci U S A 119: e2202799119.

Zhong Y, Nellimoottil T, Peace JM, Knott SR, Villwock SK, Yee JM, Jancuska JM, Rege S, Tecklenburg M, Sclafani RA et al. 2013. The level of origin firing inversely affects the rate of replication fork progression. J Cell Biol 201: 373–383.

Zhu W, Shi Y, Zhang C, Peng Y, Wan Y, Xu Y, Liu X, Han B, Zhao S, Kuang Y et al. 2023. In-frame deletion of SMC5 related with the phenotype of primordial dwarfism, chromosomal instability and insulin resistance. Clin Transl Med 13: e1007.

