## Supplementary Figures for "The SMC5/6 complex prevents genotoxicity upon APOBEC3A-mediated replication stress"

### SUPPLEMENTARY MATERIALS

#### SUPPLEMENTARY FIGURES & LEGENDS

Figure S1

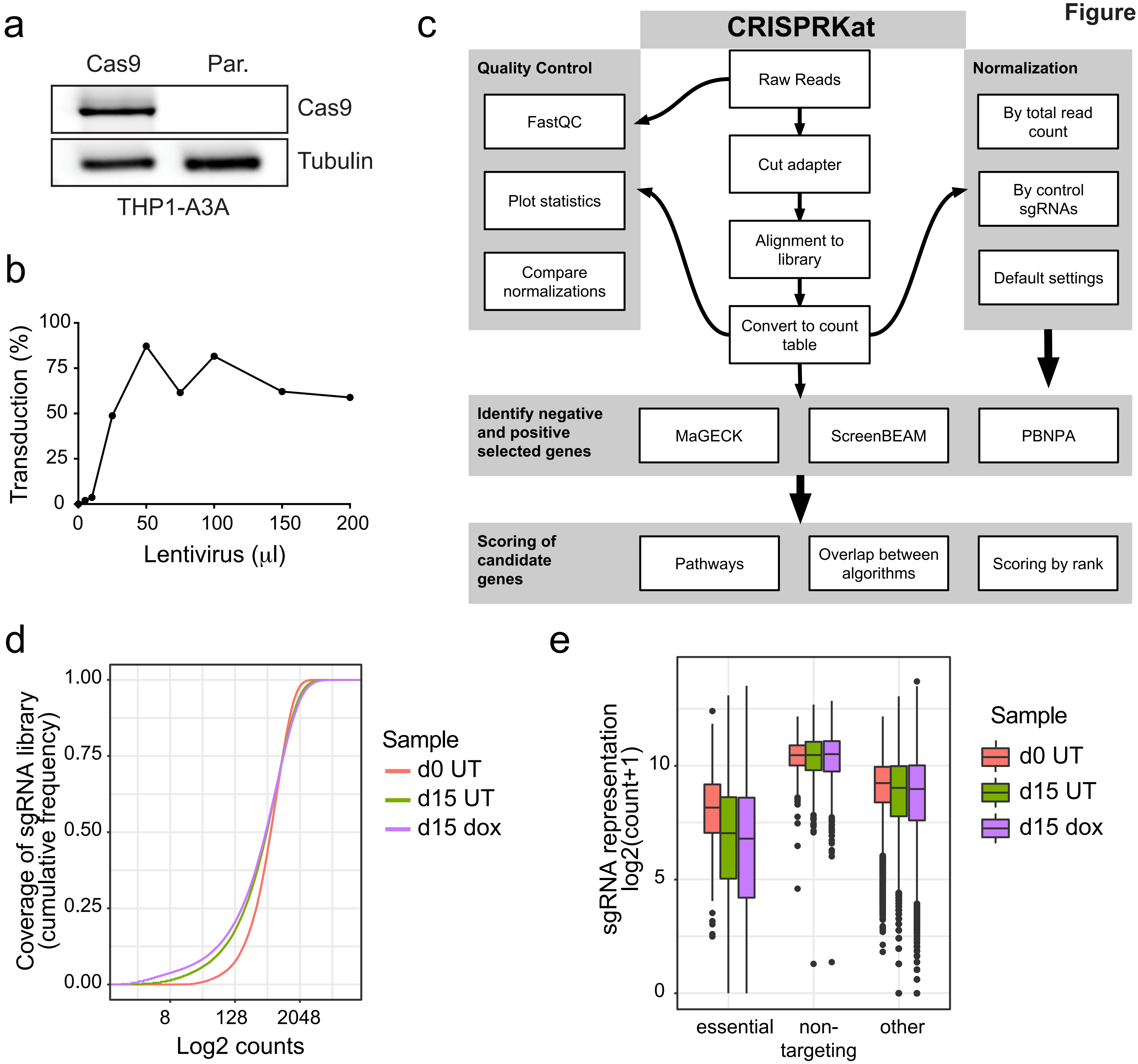

**Supplementary Figure 1. CRISPR screen design and analysis.** (**a**) A Cas9 transgene was introduced into THP1-A3A cells by lentiviral integration. Immunoblot shows Cas9 expression in transduced cells compared to parental cell line. Tubulin serves as a loading control. (**b**) The Brunello sgRNA library was integrated into THP1-A3A cells by lentiviral transduction to achieve a 1:1 ratio of lentivirus to cells to knock out one gene per cell. Two million cells were transduced with varying volumes of concentrated library lentivirus. Transduction efficiency is estimated by number of transduced cells treated with puromycin selection compared to untreated. Screen was performed with 25 µL virus/$2x10^6$ cells (see Methods). (**c**) The CRISPRKat tool flowchart shows the process from raw NGS reads to the final candidate gene list. Raw reads undergo quality control, trimming, and alignment to the Brunello sgRNA library. To control for varied sequence depth and batch effect, counts are normalized in multiple different ways as indicated and prioritized by clustering. Results are analyzed by multiple commonly used CRISPR screen analysis tools as indicated to generate a composite gene score for each sgRNA represented in the baseline sequencing from day 0. (**d**) Cumulative frequency of the normalized sgRNA counts for each condition sequenced. (**e**) Representation of essential gene sgRNAs and non-targeting control sgRNAs across samples.

Figure S2

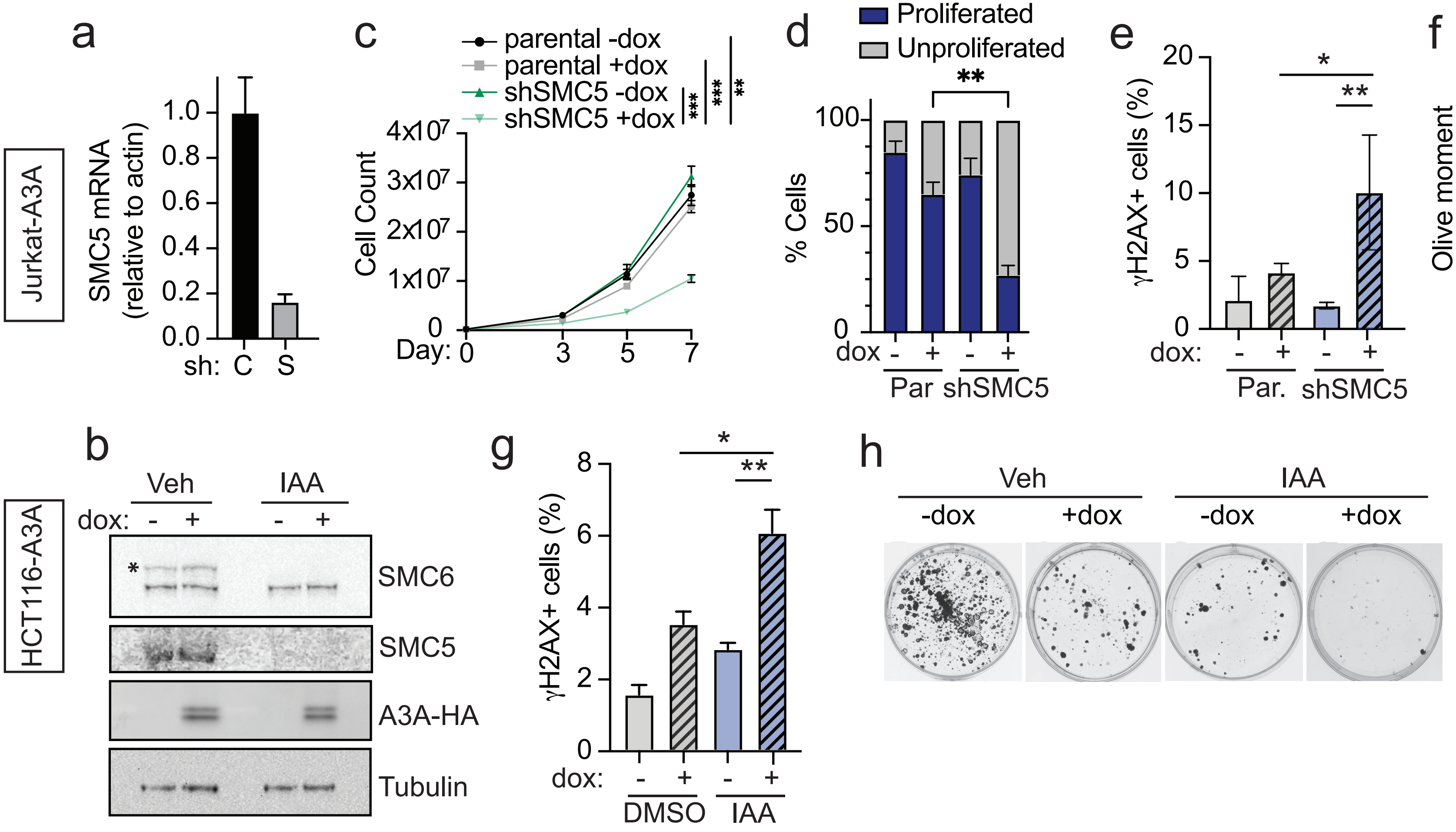

a
Jurkat-A3A
SMC5 mRNA (relative to actin)
1.0
0.8
0.6
0.4
0.2
0.0
sh: C S
c
parental -dox
parental +dox
shSMC5 -dox
shSMC5 +dox
***
***
**
Cell Count
4x10^7
3x10^7
2x10^7
1x10^7
0
Day: 0 3 5 7
d
Proliferated
Unproliferated
**
% Cells
100
50
0
dox - + - +
Par shSMC5
e
γH2AX+ cells (%)
20
15
10
5
0
*
**
dox: - + - +
Par. shSMC5
f
Olive moment
80
60
40
20
0
****
*
dox: - + - +
Par. shSMC5
b
HCT116-A3A
Veh
IAA
dox: - + - +
*
SMC6
SMC5
A3A-HA
Tubulin
g
γH2AX+ cells (%)
8
6
4
2
0
*
**
dox: - + - +
DMSO IAA
h
Veh
IAA
-dox
+dox
-dox
+dox

**Supplementary Figure 2. SMC5/6 depletion and APOBEC3A expression result in proliferative defects and DNA damage in multiple cell types.** (**a**) Jurkat cells engineered to express doxycycline-inducible HA-tagged APOBEC3A (Jurkat-A3A) were depleted of SMC5 (S) by integration of short-hairpin RNA (shRNA). SMC5 transcript quantitation is compared to Jurkat-A3A cells with integrated non-targeting shRNA control (C). n=3 technical triplicate, error bars are SD. (**b**) HCT116 cells with integrated Ostir and mAID tags on endogenous SMC6 and NSE4A loci were engineered to express dox-inducible HA-tagged APOBEC3A (HCT116-A3A). Cells were treated for 72 hours with indoleacetic acid (IAA) to degrade SMC6 and NSE4 and dox to induce APOBEC3A. Lysates from cells treated with IAA, dox, or combination are compared to untreated HCT116-A3A cells. Immunoblot was probed with antibodies targeting HA tag, SMC6, and SMC5. Asterisk (*) indicates SMC6-specific band. Tubulin was used as a loading control. (**c-f**) Jurkat-A3A cells were treated with dox every other day. (**c**) Cell proliferation was measured by counting cells over 7 days. n=3 biological replicates, error bars are SEM, p-value by sum-of-squares F-test. (**d**) Proliferation was assessed by intracellular CFSE staining, analyzed by flow cytometry. (**e**) Flow cytometry analysis of intracellular staining for $\gamma$H2AX after 72 hours of dox treatment. (**f**) Neutral comet assay to detect DSBs after 72 hours of dox treatment. For panels **d-f**, n=3 biological replicates, p-value by two-tailed t-test, error bars are SD. (**g**) HCT116-A3A cells were treated as in panel b. Flow cytometry analysis of intracellular staining for $\gamma$H2AX, n=3 biological replicates, error bars are SEM, p-value by two-tailed t-test. (**h**) HCT116-A3A cells were treated as in panel b, seeded at low density, and cultured for 2 weeks. Remaining colonies were stained with crystal violet. *p<0.05, **p<0.01, ***p<0.001, ****p<0.0001.

Figure S3

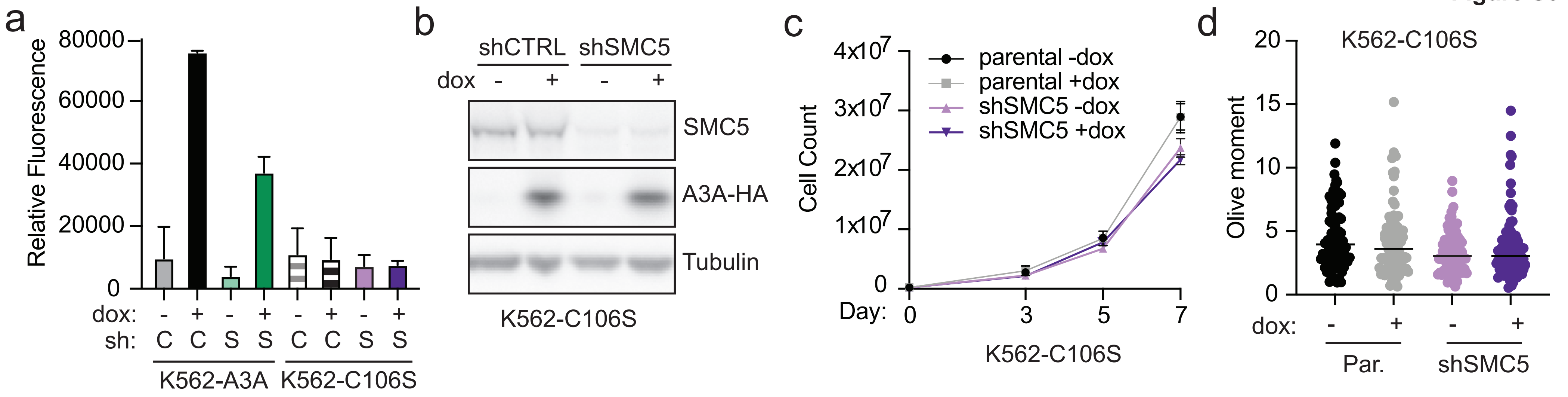
a
Relative Fluorescence
80000
60000
40000
20000
0
dox: - + - + - + - +
sh: C C S S C C S S
K562-A3A
K562-C106S
b
shCTRL
shSMC5
dox - + - +
SMC5
A3A-HA
Tubulin
K562-C106S
c
Cell Count
4x10^7
3x10^7
2x10^7
1x10^7
0
Day: 0 3 5 7
parental -dox
parental +dox
shSMC5 -dox
shSMC5 +dox
K562-C106S
d
K562-C106S
Olive moment
20
15
10
5
0
dox: - + - +
Par.
shSMC5

**Supplementary Figure 3. Synthetic lethality is dependent on deaminase activity of APOBEC3A.** (**a**) Relative fluorescence indicates C>U deamination activity in cell lysates from K562-C106S cells in comparison to K562-A3A cells. Lysates from cells with shRNA targeting control (C) or SMC5 (S). Error bars are SD. (**b**) K562 cells engineered to express dox-inducible HA-tagged A3A*C106S (K562-C106S), a catalytically inactive APOBEC3A mutant, were depleted of SMC5 by RNAi (shSMC5). Immunoblot shows A3A*C106S expression after 24 hours of dox induction. Blot was probed with antibodies targeting HA tag and SMC5. Tubulin was used as a loading control. (**c**) Cell proliferation was measured by counting cells over 7 days. Error bars are SEM. p-value by sum-of-squares F test was not significant for all comparisons. (**d**) Neutral comet assay to detect DSBs after 72 hours of dox treatment. Bar is median, error bars are SD. No comparisons are significant by two-tailed t-test. All assays represent results of three independent biological replicates.

**Figure S4**

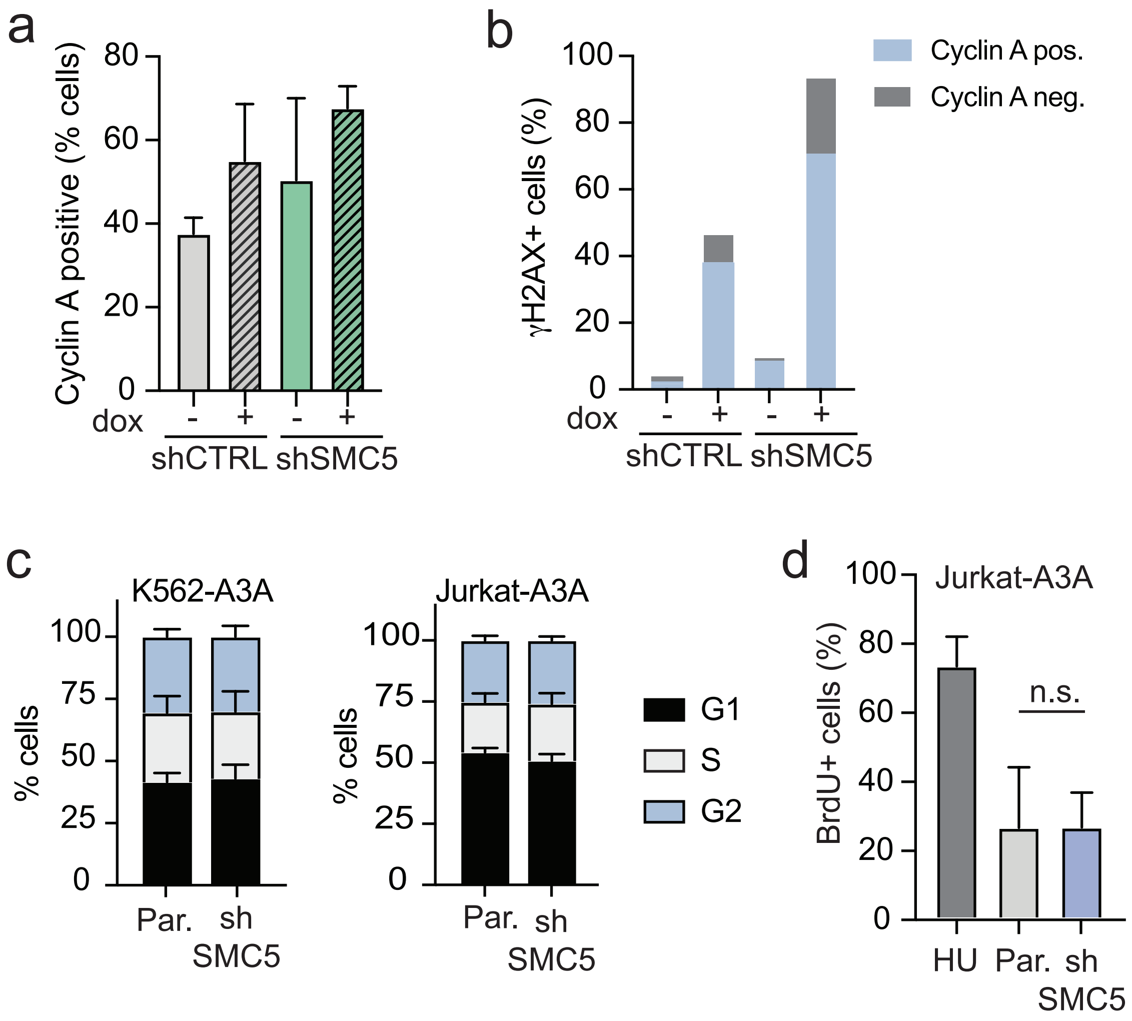
a
Cyclin A positive (% cells)
80
60
40
20
0
dox
-
+
-
+
shCTRL
shSMC5
b
γH2AX+ cells (%)
100
80
60
40
20
0
dox
-
+
-
+
shCTRL
shSMC5
Cyclin A pos.
Cyclin A neg.
c
K562-A3A
Jurkat-A3A
% cells
100
75
50
25
0
Par.
sh SMC5
G1
S
G2
d
Jurkat-A3A
BrdU+ cells (%)
100
80
60
40
20
0
n.s.
HU
Par.
sh SMC5

**Supplementary Figure 4. SMC5/6 depletion does not alter cell cycle or increase ssDNA accumulation.** K562-A3A cells were treated, stained, and imaged as in Figure 5. (**a**) Quantification of cyclin A-positive cells. Error bars are SEM. (**b**) Quantification of all cells that had $\geq$5 nuclear γH2AX foci, divided by cyclin A-positive or negative staining. Bars are mean, n=3 biological replicates. At least 200 nuclei analyzed per condition. (**c**) K562-A3A and Jurkat-A3A cells were depleted of SMC5 by RNAi, stained with propidium iodide, and analyzed for cell cycle profile by FACS. n=3 biological replicates, bars are mean with SD. (**d**) Native BrdU staining was performed on Jurkat-A3A cells. Cells were pulsed with BrdU for 36 hours. Intracellular staining with anti-BrdU antibody was analyzed by flow cytometry. P-value by two-tailed t-test, n=3 biological replicates, bars are mean with SD. HU-treated Jurkat cells were used as positive control.

**Figure S5**

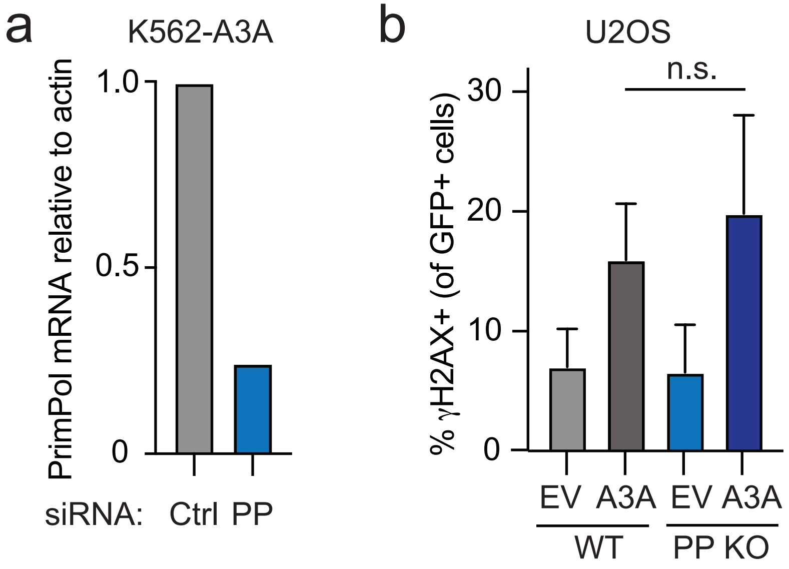
a
K562-A3A
PrimPol mRNA relative to actin
1.0
0.5
0
siRNA:
Ctrl
PP
b
U2OS
n.s.
% γH2AX+ (of GFP+ cells)
30
20
10
0
EV
A3A
EV
A3A
WT
PP KO

**Supplementary Figure 5. PrimPol depletion does not exacerbate APOBEC3A-mediated DNA damage.** (**a**) Quantitative PCR demonstrates PrimPol depletion in K562-A3A cells by siRNA transfection. All PrimPol mRNA levels displayed relative to siCtrl in parental cells. (**b**) U2OS cells (wild-type or PrimPol KO) were transfected with empty vector (EV) or APOBEC3A-containing plasmid and analyzed for γH2AX by flow cytometry after 72 hours. All vectors express GFP, thus the fraction of γH2AX within GFP+ cell populations which reflect only transfected cells is shown. Bars are mean with SD, n=3 biological replicates, p-value by two-tailed t-test.

#### SUPPLEMENTARY MATERIALS AND METHODS.

##### Genome-wide CRISPR-Cas9 knockout screen in THP1-A3A cells

*Brunello library lentivirus titration*. Pooled lentivirus encoding the Brunello guide RNA library was generated as previously described[82]. To find optimal transduction conditions for an MOI of 0.3-0.5, cells were transduced with the Brunello library via spinfecting $2x10^6$ cells with a range of virus volumes. Cells were seeded in a 12-well plate at $2x10^6$ cells per well in 1 mL standard media supplemented with 10 µg/mL polybrene (Santa Cruz). Lentivirus was placed in the wells, along with a no-transduction control. The 12-well plate was centrifuged 2,000 rpm for 90 minutes at room temperature and incubated overnight.

After overnight incubation, cells were diluted 1:10 and 1:20 into four replicate plates. One plate for each dilution received 1 µg/mL puromycin. After 3 days, cells were counted to calculate transduction efficiency. Percent transduction was calculated by cell count from the dilution replicate with puromycin divided by cell count from the dilution replicate without puromycin multiplied to 100. The virus volume yielding a MOI closest to 0.4 was chosen for large-scale screening.

*Genome-Wide Screen Analysis*. Using the guide sequencing data, we analyzed enrichment and depletion of gRNAs using a multi-modality algorithm that we named CRISPRKat (**Fig S1C**) after its inventor, Katharina Hayer. Features of CRISPRKat include incorporation of several quality control measures, multiple normalization methods, and input from a variety of publicly available CRISPR screen analysis tools [Model-based Analysis of Genome-wide CRISPR/Cas9 Knockout (MAGeCK), Permutation-Based Non-Parametric Analysis (PBNPA), and Screening Bayesian Evaluation and Analysis Method (ScreenBEAM) algorithms] to identify negatively selected genes. Candidate genes are further weighted by biological variables, ultimately generating a novel scoring system used to rank genes.

**Native BrdU detection.** Cells were plated at a density of $5x10^5$ cells per well in a 6-well plate. After one population doubling, cells were pulsed with 10 µL 10 µM BrdU (BD Biosciences) per well. After two additional population doublings, cells were harvested, pre-extracted with 1 mL PBS with 0.2% Triton-X, and then the BD Biosciences FITC BrdU Flow Kit staining protocol was followed beginning from Step 2.

**Deamination assay.** K562 cells were treated with dox to induce expression of A3A or C106S as described. Cell pellets were collected by centrifugation. Cells were lysed on ice for 10 minutes with 1X RIPA buffer (Cell Signaling Technology) with the addition of 1X Pierce protease inhibitor (EDTA Free, Thermo Scientific) and PMSF (GoldBio). Lysate was sonicated briefly and pelleted by centrifugation. The supernatant was quantified using Coomassie reagent and BSA standards (Thermo Scientific) and a POLARstar Omega plate reader (BMG LABTECH). A deaminase reaction buffer containing 50mM tris HCl pH 7.4 and EDTA pH 8 was prepared in water and the pH was adjusted to 5.9-6.1. After pH adjustment, uracil DNA glycosylase (NEB) was added to the buffer (2.5U/sample) to induce excision of uracil bases. For each sample, 5µg of lysate was combined with the reaction buffer and an oligo containing a single cytosine base (AAATTCAGAGAGAGAATGTGA) with a 56-FAM fluorophore and 36-TAMSp quencher (IDT) in a flat-bottom, clear 96-well plate. Control reactions contained RIPA buffer incubated with either the cytosine oligo (56-FAMAAATTCAGAGAGAGAATGTGA36-

TAMSp, negative control) or an identical oligo with uracil in place of the cytosine (56-FAMAAATTUAGAGAGAGAATGTGA36-TAMSp, positive control, not shown). All samples and control reactions were performed in technical triplicate. The reaction mixture was incubated at 37°C for 1.5 hours, followed by the addition of 3uL of 4N NaOH for 30 minutes at 37°C to induce cleavage of the oligo at any resulting abasic sites. Oligo cleavage separates the fluorophore and quencher resulting in fluorescence. Samples were neutralized with 3uL 4N HCl and 27uL 2M Tris pH 7.9 per sample. The 96-well plate was cooled at 4°C for several minutes before analysis. Fluorescence was measured using a POLARstar Omega plate reader (BMG LABTECH) with excitation set to 485nm and emission set to 520nm to detect the FAM fluorophore. The technical triplicate values were averaged, and the background fluorescence (cytosine-containing oligo with RIPA buffer) was subtracted from each sample. The graph shows the combined results of three biological replicates.
